# Functional and multi-omic aging rejuvenation with GLP-1R agonism

**DOI:** 10.1101/2024.05.06.592653

**Authors:** Junzhe Huang, Andrew J. Kwok, Jason Chak Yan Li, Clement Lek Hin Chiu, Bonaventure Y. Ip, Lok Yi Tung, Xianyi Zheng, Hoi Tung Chow, Michelle P. S. Lo, Zhongqi Li, Roy C. H. Chan, Nenghan Lin, Ziyu Wang, Manyu Wang, Leo Y. C. Yan, Danny C. W. Chan, William K. K. Wu, Kim Hei-Man Chow, Wei-Jye Lin, Yamei Tang, Billy Wai-Lung Ng, Sunny H. Wong, Thomas W. Leung, Vincent C. T. Mok, Ho Ko

## Abstract

Identifying readily implementable methods that can effectively counteract aging is urgently needed for tackling age-related degenerative disorders. Here, we conducted functional assessments and deep molecular phenotyping in the aging mouse to demonstrate that glucagon-like peptide-1 receptor agonist (GLP-1RA) treatment attenuates body-wide age-related changes. Apart from improvements in physical and cognitive performance, the age-counteracting effects are prominently evident at multiple omic levels. These span the transcriptomes and DNA methylomes of various tissues, organs and circulating white blood cells, as well as the plasma metabolome. Importantly, the beneficial effects are specific to aged mice, not young adults, and are achieved with a low dosage of GLP-1RA which has a negligible impact on food consumption and body weight. The molecular rejuvenation effects exhibit organ-specific characteristics, which are generally heavily dependent on hypothalamic GLP-1R. We benchmarked the GLP-1RA age-counteracting effects against those of mTOR inhibition, a well-established anti-aging intervention, observing a strong resemblance across the two strategies. Our findings have broad implications for understanding the mechanistic basis of the clinically observed pleiotropic effects of GLP-1RAs, the design of intervention trials for age-related diseases, and the development of anti-aging-based therapeutics.

## Main

Aging is a complex process involving diverse cellular and molecular alterations across all body systems, resulting in progressive functional decline. Discovering effective strategies to counteract aging-associated changes is a significant scientific pursuit with profound societal implications, given the potential to improve overall well-being and extend healthy lifespan. Numerous anti-aging strategies have shown promising experimental data, including mammalian target of rapamycin (mTOR) inhibitors^1,2^, senolytics^3,4^, nicotinamide adenine dinucleotide boosters^5–7^, taurine supplements^8^, intermittent fasting and calorie restriction^9–11^, cellular reprogramming^12,13^, and circulating factor or protein-based rejuvenation^14–22^. Studies testing these methods have significantly advanced our understanding of the aging process and enhanced our ability to counteract it. An ideal rejuvenation method should possess several characteristics, including: (*i*) a pharmacological approach to facilitate practical deployment, (*ii*) a good safety profile with a wide therapeutic window for ease of achieving a good benefit-side effect balance, (*iii*) broad potential applicability to various diseases affecting different organ systems in aging, and (*iv*) potential mechanistic synergy with other therapeutic targets in age-related diseases, allowing for combination therapies. While the above approaches are promising, each has its own set of challenges and limitations. To date, many still require further refinement and development in one or more key attributes for clinical application^23^.

Glucagon-like peptide-1 (GLP-1) is a peptide hormone produced by intestinal enteroendocrine cells in the periphery, and by preproglucagon-expressing neurons in the brainstem solitary tract nucleus (NTS) in the central nervous system (CNS)^24^. In the pancreas, GLP-1 receptor (GLP-1R) signaling enhances postprandial and hyperglycemia-induced insulin release. In the CNS, GLP-1R in the hypothalamus and the brainstem NTS play crucial roles in regulating satiety, metabolism, and other neuroendocrine processes^24–26^. With advancements in understanding GLP-1 biology, multiple GLP-1R agonists (GLP-1RAs) have been developed based on pharmacokinetic innovations, and have achieved remarkable success in treating diabetes mellitus (DM) and obesity^27^. Notably, besides cardiovascular and renal benefits^28–31^, GLP-1RA use in DM has shown a wide range of pleiotropic effects, including a reduction in the incidence of cognitive decline^32,33^, Parkinson’s disease^34^, and cancers of some organs^35–37^. In non-diabetic overweight or obese subjects, use of GLP-1RA reduced cardiovascular mortality^38^. GLP-1RAs have also demonstrated efficacy in various animal models of neurodegenerative conditions^39,40^. Pilot trials investigating the use of GLP-1RAs in the treatment of non-diabetic patients with dementia^41,42^ or Parkinson’s disease^43,44^ have been conducted, with encouraging results.

We postulate that GLP-1RAs are able to broadly counteract age-related changes body-wide. This would offer a unifying mechanistic explanation for the drugs’ effectiveness in diverse disease models and their expanding clinical indications. As a pharmacological approach, GLP-1R agonism also fulfills the above-mentioned criteria for an ideal rejuvenation method. We have previously demonstrated that treatment with exenatide (aka exendin-4, a GLP-1RA) potently counteracts age-related transcriptomic changes across diverse glial and vascular cell types in the mouse brain^45,46^. In this study, we investigate whether GLP-1RA treatment can impart whole-body rejuvenation effects. Such effects, if present, should span across organs, lead to functional improvements, and manifest at several molecular levels, including the transcriptome, DNA methylome, and the circulating metabolome. As GLP-1R is highly expressed by subsets of cells in the hypothalamus^47^, a key regulator of systemic homeostatic processes^48^, we also aim to determine whether the molecular age-counteracting effects depend on CNS GLP-1 signaling through hypothalamic GLP-1R. Finally, we benchmark our GLP-1RA strategy against the mTOR inhibitor rapamycin – currently the most potent pharmacological anti-aging agent.

### GLP-1R agonist treatment improves physical and cognitive functions in aging mice

We first asked if long-term GLP-1RA treatment can improve physical and cognitive functions in aged mice. To this end, we administered male C57BL/6 mice with either intraperitoneal injection (I.P.) of a GLP-1RA (exenatide, 5 nmol/kg bw/day, *n* = 9 animals) or a vehicle (PBS, *n* = 9 animals) for a duration of 30 weeks, starting when the mice were 11 months old (m.o.), and conducted functional assessments on these animals (**Fig. 1a**). To evaluate the impacts on muscle power and motor coordination, we carried out forelimb grip strength and accelerated rotarod tests at baseline, after 3 months, and after 6 months of treatment. Additionally, we employed the Barnes maze, a spatial learning and memory assay, to assess cognitive performance after 6 months of treatment. The exenatide-treated mice demonstrated a progressive enhancement in both forelimb grip strength (**Fig. 1b**) and rotarod test performances (**Fig. 1c**) relative to the vehicle group animals of the same age and treatment duration. The effect was more pronounced after 6 months of treatment compared to 3 months (**Fig. 1b, c**). Notably, the exenatide-treated group also exhibited a significant advantage in the Barnes maze test after 6 months of treatment (**Fig. 1d**). These mice learnt to reach the target more quickly compared to the vehicle control group, which took four days to achieve a comparable performance (**Fig. 1d**).

**Figure 1.**
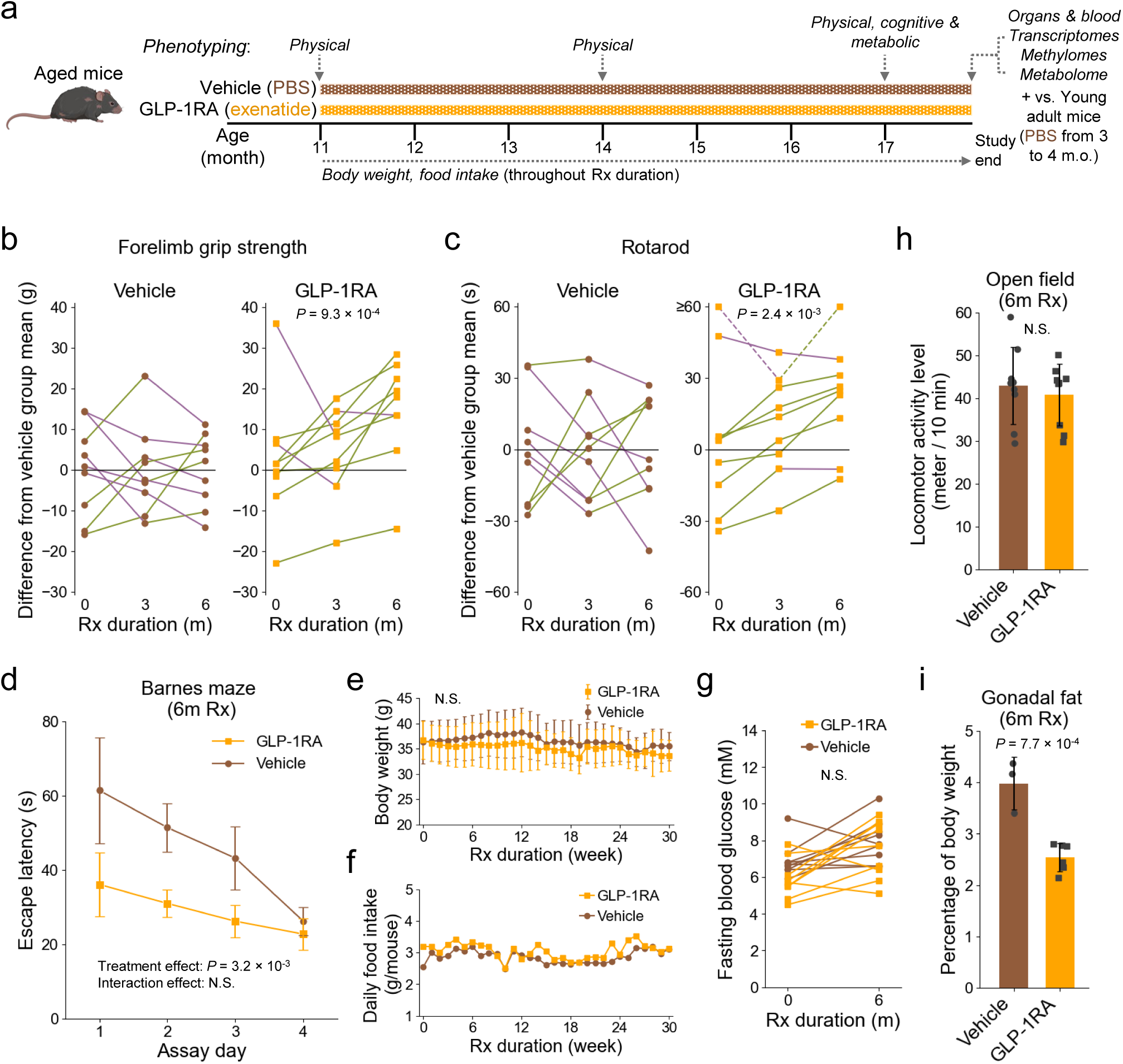
Physical, cognitive and metabolic readouts in aging mice treated with a GLP-1RA (exenatide) for 30 weeks starting at 11 months old. a, Schematic of experimental design for the aged long-term treatment cohort. b, The relative performances of same-age mice treated with exenatide vs. phosphate-buffered solution (PBS) vehicle for the same durations in forelimb grip strength test. Values are plotted as differences from vehicle group mean at each treatment duration time point. Dots connected by lines are longitudinal data from individual animal subjects. The lines are color-coded according to temporal change across two consecutive time points of assessment (green: increase; magenta: decrease). c, Similar to b, but for performances in the accelerated rotarod test, reflecting motor coordination ability. Dashed lines in the right panel indicate lines connecting to dots with values beyond the *y*-axis upper limit. d, Comparison of the performances of exenatide and vehicle-treated mice in the Barnes maze test, a spatial memory-dependent task, after 6-month treatment. Mean (±S.E.M.) time to escape for animals over four consecutive assay days are shown for each treatment group. Shorter times indicate better performances. e, Mean (±S.D.) body weight of the two animal groups throughout the treatment period (monitored weekly). f, Average daily food intake per mouse for the two animal groups throughout the treatment period (monitored weekly). g, Fasting blood glucose levels at baseline and after 6-month treatments for each animal subject in the two groups. h, Comparison of locomotor activity during a 10-minute open field test after 6-month treatments. i, Gonadal fat weight as percentage of body weight in the two treatment groups. Sample sizes: *n* = 9 mice for each experimental group for data presented in b–h, and *n* = 3 and 6 mice for exenatide and vehicle treatment groups respectively for data presented in i. *P*-values: b and c, Page’s trend test; d, e and g, two-way repeated measures ANOVA; h and i, two-sided unpaired *t*-test. N.S. in d, e, g, and h indicate statistical non-significance with *P*-value > 0.05. Abbreviation: Rx, treatment.

To control for potential confounding factors that could impact aging, we closely monitored body weight and food intake throughout the treatment period, while fasting blood glucose and locomotor activity level in an open field assay were also assessed after 6-month treatment. Importantly, at the chosen dosage of exenatide, there were no significant differences in body weight (**Fig. 1e**) or food intake (**Fig. 1f**) between the two groups during the treatment period. Additionally, there were no observed differences in fasting blood glucose (**Fig. 1g**) or physical activity (**Fig. 1h**) levels between the two groups after 6 months of treatment. The exenatide-treated group exhibited a reduction in gonadal fat as a percentage of body weight (**Fig. 1i**), which aligns with the known pharmacological action of GLP-1R agonism^49^.

We also carried out a separate set of experiments in young male mice (*n* = 9 animals each group), starting at 3 m.o. (**Supplementary Fig. 1a**). In contrast to the aged mice, the young mice treated with exenatide did not perform better in the forelimb grip strength test (**Supplementary Fig. 1b**), and only exhibited a marginal improvement in the rotarod test (**Supplementary Fig. 1c**). The young exenatide-treated animals also did not outperform the vehicle-treated mice in Barnes maze (**Supplementary Fig. 1d**), and displayed a slight increase in overall locomotion in the open field test (**Supplementary Fig. 1e**) after 6 months of treatment. Exenatide reduced body weight gain in the young mice (**Supplementary Fig. 1f**), despite their food intake remaining unaffected (**Supplementary Fig. 1g**). Similar to the aged animals, there were no observed differences in fasting blood glucose between the two groups after 6 months of treatment (**Supplementary Fig. 1h**), whereas gonadal fat was also reduced by the long-term exenatide treatment (**Supplementary Fig. 1i**).

Collectively, these results indicate that GLP-1RA treatment significantly improves physical and cognitive functions in aged mice. The observed effects can be achieved at a dosage that has minimal impact on metabolic measures, and they are predominantly noticeable in aged animals. This suggests that the beneficial impact of GLP-1RA treatment is particularly pronounced in the context of aging.

### GLP-1R agonism ameliorates body-wide age-related molecular changes across multiple omic levels

We next aimed to explore whether the observed functional benefits were associated with the counteraction of age-related molecular changes. To this end, at the conclusion of the treatment period we collected various tissue organs and blood samples from the animals for multi-omic assessments (**Fig. 1a**). A young control group that received vehicle treatment from 3 months of age for 4 weeks was additionally included for comparative analysis (**Fig. 1a**).

Using bulk RNA sequencing, we compared the age-related and exenatide treatment-induced transcriptomic changes throughout the body. Strikingly, exenatide treatment led to a widespread, global counteraction of age-related transcript expression changes in numerous tissue organs, as well as in circulating white blood cells (WBCs) (**Fig. 2a–h, Supplementary Fig. 2a–e**). This effect was particularly prominent in metabolically demanding tissues and organs, including the hypothalamus, hippocampus, frontal cortex (**Fig. 2a, b, Supplementary Fig. 2a**), gonadal adipose tissue (**Fig. 2c**), colon (**Fig. 2d**), heart (**Fig. 2f**), and skeletal muscle (**Fig. 2g**). The age-related transcriptomic changes that were counteracted by exenatide treatment varied across the different tissue organs and circulating WBCs (**Fig. 2a–g, right panels**), consistent with the known organ-specific patterns of aging-associated expression changes^50^. Additionally, exenatide treatment counteracted age-related metabolomic changes in the circulation (**Fig. 2i**), apart from the effect observed in the circulating WBC transcriptome (**Fig. 2e**).

**Figure 2.**
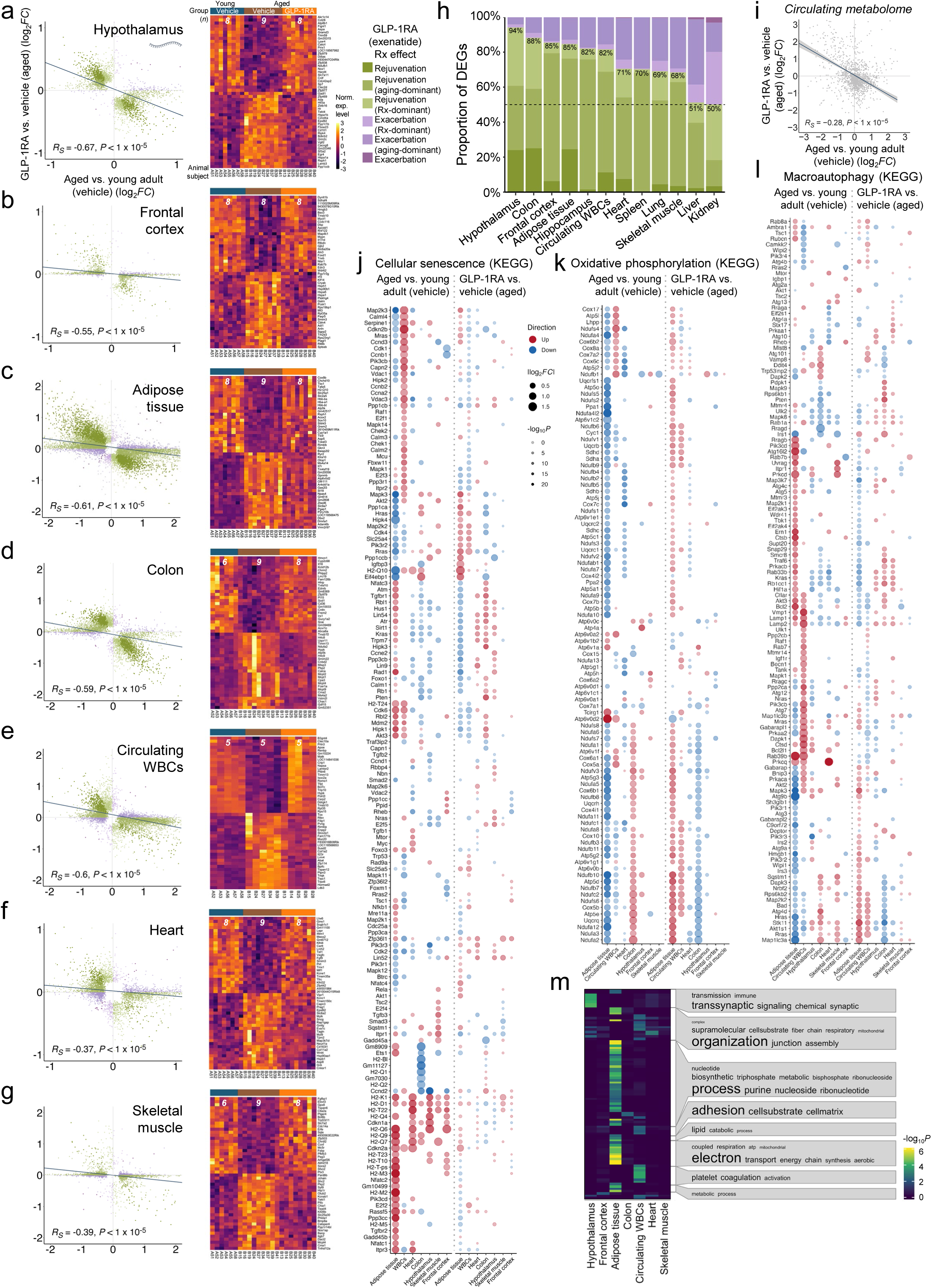
Transcriptomic impacts across tissue organs and circulating white blood cells (WBCs) in aging mice treated with a GLP-1RA (exenatide) for 30 weeks starting at 11 months old, and associated metabolomic changes. **a**–**g**, Left panels: Scatter plots showing transcriptomic changes in aging (*x*-axis) vs. exenatide treatment (*y*-axis) in different tissue organs and circulating WBCs. Each dot represents one differentially expressed gene (DEG), color-coded by treatment (Rx) effect categories as shown in the inset (rejuvenation: opposing aging and Rx effects; exacerbation: same aging and Rx effects; aging- or Rx-dominant: statistically significant in the respective comparison only; no additional label: significant for both aging and Rx effect comparisons, also see **Methods**). Inset of **a**: symbol for transcript. Right panels: Heatmaps of the expression levels of top 50 protein-coding genes by log2 fold change with aging-associated differential expressions (25 up- and 25 down-regulated) that were reversed by exenatide treatment in the different tissue organs and circulating WBCs. Number at the top of each heatmap: samples sizes (numbers of mice) for each tissue. **h**, Summary bar plot of the proportions of DEGs under the different Rx effect categories (as specified in the inset of **a**). **i**, Scatter plot showing plasma metabolomic changes in aging (*x*-axis) vs. exenatide treatment (*y*-axis). Each dot represents one metabolite. **j**–**l**, Dot plots showing DEGs annotated with three aging-associated functional pathways (KEGG database). Genes with significant differential expression in at least one comparison (i.e., aging and/or treatment effects) are included. **m**, Functional pathway terms enrichment among aging-associated transcripts with differential expressions that were rejuvenated by exenatide treatment across the different tissue organs and circulating WBCs. In **a**–**g** and **i**, the lines represent linear fits (with confidence interval (grey) in **i**). The Spearman correlation coefficients (*R_S_*) and associated *P*-values are also shown.

Many functionally relevant genes, including those associated with key hallmarks of aging^51,52^, such as cellular senescence (**Fig. 2j**), oxidative phosphorylation (**Fig. 2k**), and macroautophagy (**Fig. 2l**), exhibited predominantly opposite patterns of transcript expression level changes between aging and exenatide treatment across the tissue organs and circulating WBCs. Enrichment analysis on the age-related transcript expression changes rejuvenated by exenatide treatment implicated diverse impacts on organ/tissue-specific functions, as well as shared processes across body sites (**Fig. 2m, Supplementary Fig. 2f**). Weighted gene co-expression network analysis^53^ (WGCNA) also demonstrated that exenatide treatment modulated the expression of gene modules in a manner counteractive to aging across the tissue organs and circulating WBCs (**Supplementary Fig. 3a**). Notably, while the differentially expressed modules differ across the tissue organs (**Supplementary Fig. 3a**), many of them were enriched in genes involved in biological processes that undergo age-dependent changes, such as autophagy/mitophagy, cellular senescence, and proteostasis (**Supplementary Fig. 3b, c**).

Epigenetic changes play a crucial role in regulating gene expression and are recognized as a hallmark of aging^51,52^. We further investigated the effects of GLP-1RA treatment on DNA methylation patterns and their correlation with age-related modifications, using a DNA microarray that covers over 320 thousand CpG sites^54,55^. These consist of ∼285 thousand mouse CpG sites^56^, and also allows imputation of ∼40 thousand sites conserved among multiple mammalian species, the effects on which may have higher generalizability across species^54,55^. We first analyzed the treatment impact on the ∼285 thousand mouse sites. Similar to the transcriptomic level, we found that exenatide treatment effectively and globally counteracted aging-associated methylation changes across multiple tissues, encompassing brain regions (the hypothalamus, hippocampus, and frontal cortex), gonadal adipose tissue, circulating WBCs, heart, and skeletal muscle (**Fig. 3a, b**). Interestingly, the rejuvenation effect was not evident on the colon or spleen methylome (**Fig. 3a, b**), despite the transcriptomic rejuvenation in the colon (**Fig. 2d**). By contrast, exenatide treatment induced methylation changes antagonistic to those that occurred in aging in the kidneys and liver (**Fig. 3a, b**), whereas we did not detect dominant transcriptomic rejuvenating effects for these organs (**Fig. 2h, Supplementary Fig. 2d, e**). We obtained similar results with additional analysis focusing on the ∼40 thousand imputed multi-mammalian conserved sites (**Fig. 3c, Supplementary Fig. 4**).

**Figure 3.**
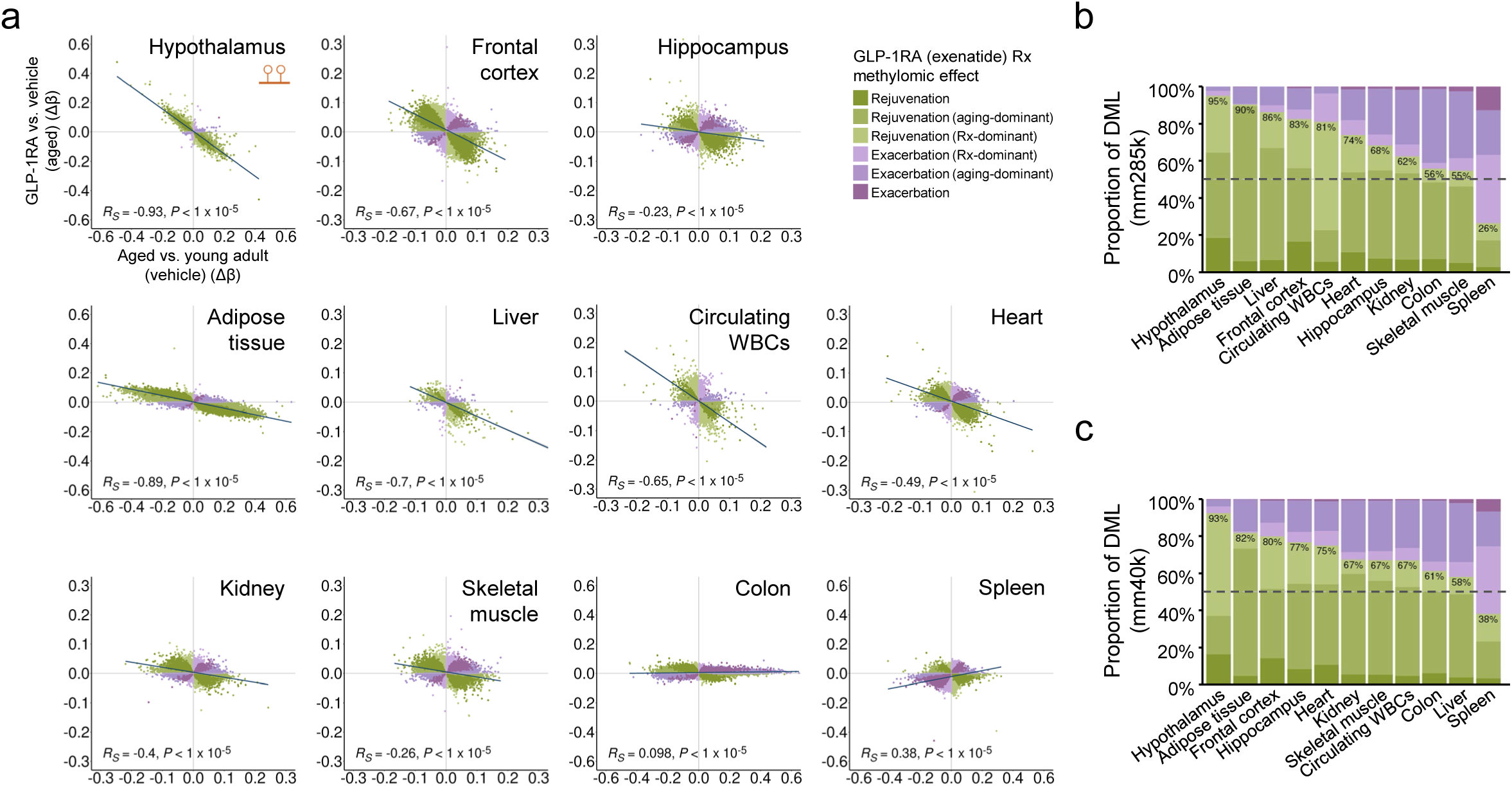
GLP-1RA (exenatide) treatment-induced methylomic changes in aging mice. **a**, Scatter plots showing DNA methylation level changes in aging (*x*-axis) vs. exenatide treatment (*y*-axis) in sites covered by the Illumina mouse 285k array (mm285k) across different tissue organs and circulating white blood cells (WBCs). Each dot represents one differentially methylated locus (DML), color-coded by treatment (Rx) effect categories as shown in the inset (rejuvenation: opposing aging and Rx effects; exacerbation: same aging and Rx effects; aging- or Rx-dominant: statistically significant in the respective comparison only; no additional label: significant for both aging and Rx effect comparisons, also see **Methods**). For each plot, the line represents linear fit. The Spearman correlation coefficients (*R_S_*) and associated *P*-values are also shown. Inset of left upper panel: symbol for DNA methylation. **b**, Summary bar plot of the proportions of DML among the mm285k array sites under different categories (as specified in the inset of **a**). **c**, Similar to **b**, but for DML among the imputed mammalian conserved sites covered by the mammalian 40k array (mm40k). Sample sizes for the various tissue organs (young adult vehicle, aged vehicle, aged exenatide): hypothalamus (4, 4, 5), frontal cortex (8, 8, 8), hippocampus (8, 8, 8), adipose tissue (8, 8, 8), liver (8, 8, 8), circulating WBCs (5, 9, 6), heart (8, 8, 8), kidney (7, 9, 8), skeletal muscle (8, 8, 8), colon (7, 9, 8), spleen (8, 8, 8).

For the young male mouse cohort (**Supplementary Fig. 1a**), we similarly profiled transcript expression changes across tissues and organs and compared the exenatide treatment-induced differential expressions to those found in the aged animals. Generally, exenatide treatment resulted in fewer and distinct expression changes in the young cohort. This was reflected by the much smaller numbers of significantly differentially expressed genes caused by exenatide treatment and the lack of correlation in exenatide induced differential expression between young and aged animals (**Supplementary Fig. 5**), except for circulating WBCs, where a modest positive correlation was observed (**Supplementary Fig. 5**). These findings indicate that the age-counteracting transcriptomic effects of GLP-1R agonist treatment are largely specific to the aged mouse.

Taken together, these findings provided compelling evidence that the functional benefits observed in aged mice following long-term GLP-1RA treatment are closely linked to potent rejuvenating effects at the transcriptomic, epigenetic, and metabolomic levels.

### Age-counteracting molecular effects of GLP-1RA depend on hypothalamic GLP-1R

The functional and multi-omic rejuvenation effects of GLP-1RA treatment can in theory be mediated by systemic impacts driven by GLP-1R activation in the CNS, the pancreas, and/or local actions at different body sites. Although it is difficult to completely separate and quantify the relative contributions of each component in every tissue organ, we recognized the importance of gaining a better understanding of the mechanisms involved in the age-counteracting effects of GLP-1RA treatment. To address this, we sought to investigate the potential role of hypothalamic GLP-1R in the age-counteracting molecular effects of GLP-1RA treatment.

We performed experiments using separate groups of male mice at an older age than the previous batch, starting treatments at 18 m.o. and continued for ∼3 months (**Fig. 4a**). These mice were divided into different groups: one group received a hypothalamic AAV injection to express a short hairpin RNA (shRNA) and specifically knock down *Glp1r* in the hypothalamus, while another group received a control AAV injection with a scramble shRNA (**Fig. 4a**). Both groups were further subdivided into subgroups that either received exenatide treatment (at the same dose as before) or PBS vehicle (**Fig. 4a**). Additionally, a young age vehicle control group that received scramble shRNA AAV injection was included for comparisons (**Fig. 4a**). To confirm the effectiveness of *Glp1r* knockdown, we assayed the transcript abundance in the hypothalamus and observed a reduction of near or over 50% in the knockdown group (**Fig. 4b**). Throughout the treatment period, food intake remained stable in these aged mice (**Supplementary Fig. 6a**), while a slight trend of decreased body weight was observed only in the exenatide-treated scramble shRNA control group (**Supplementary Fig. 6b**). No differences in glucose tolerance were found among the groups after the treatment period (**Supplementary Fig. 6c**). Although not statistically significant, the changes in gonadal fat weight were consistent with the expected effects of *Glp1r* knockdown (i.e., higher percentage of body weight with hypothalamic *Glp1r* knockdown) and GLP-1RA treatment (i.e., lower with exenatide treatment) (**Supplementary Fig. 6d**).

**Figure 4.**
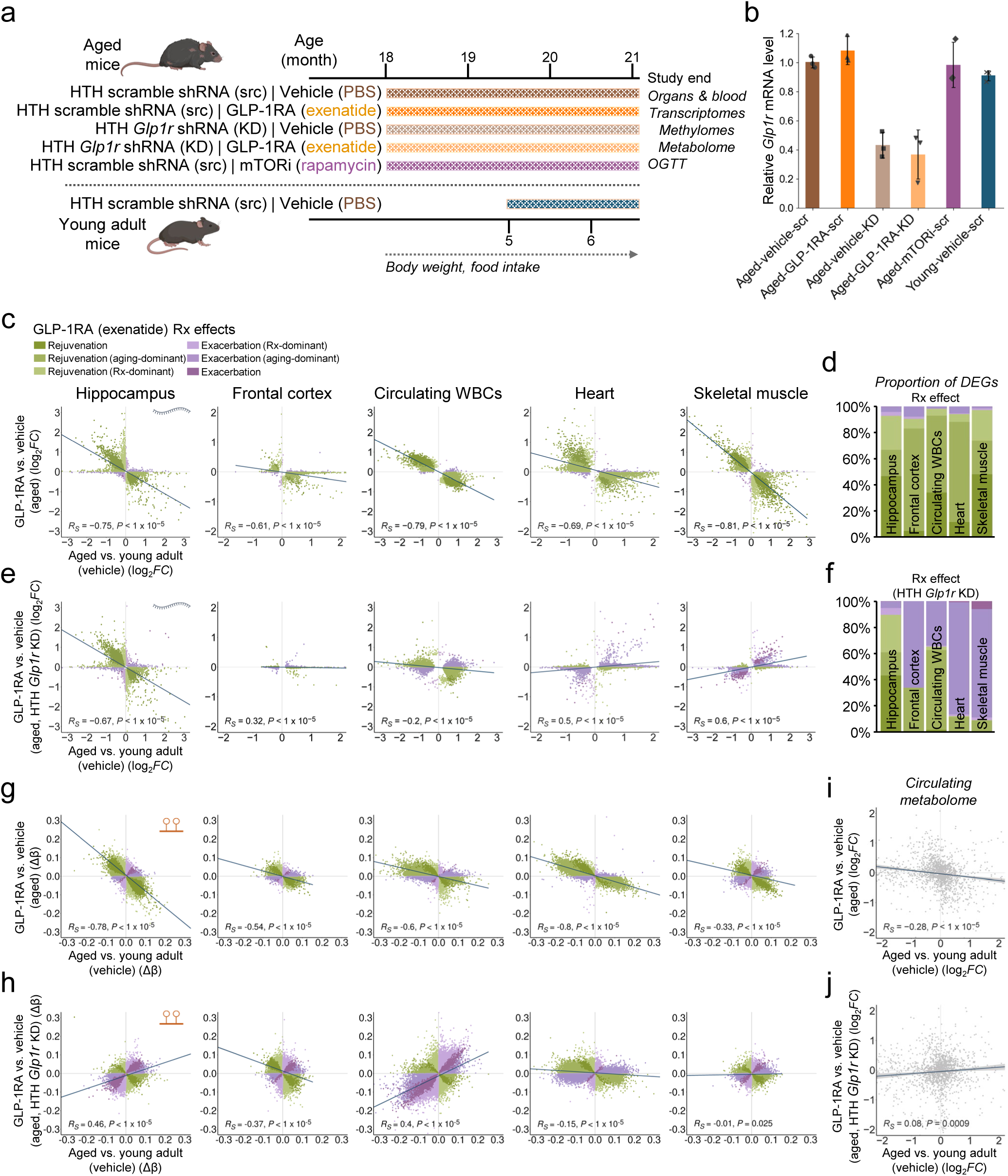
Age-counteracting transcriptomic, methylomic, and metabolomic effects of 13-week GLP-1RA (exenatide) treatment starting at 18 months old, and the dependence on hypothalamic GLP-1 receptor. **a**, Schematic of experimental design for the aged short-term treatment cohort. The animals either received hypothalamic injection of adeno-associated virus vector (AAV) for the expression of shRNA to knockdown *Glp1r* or scramble shRNA, and either received vehicle, exenatide, or rapamycin treatment. Abbreviations: HTH, hypothalamus. KD, knockdown. OGTT: oral glucose tolerance test. **b**, Mean (±S.D.) hypothalamic *Glp1r* transcript expression levels measured with quantitative PCR (*n* = 3 mice from each group used for the comparison), relative to the mean of aged control group (i.e., hypothalamic scramble shRNA-AAV injection, treated with vehicle). **c**, Scatter plots showing transcriptomic changes in aging (*x*-axis) vs. exenatide treatment (*y*-axis) in different tissue organs and circulating white blood cells (WBCs). Each dot represents one differentially expressed gene (DEG), color-coded by treatment (Rx) effect categories as shown in the inset (rejuvenation: opposing aging and Rx effects; exacerbation: same aging and Rx effects; aging- or Rx-dominant: statistically significant in the respective comparison only; no additional label: significant for both aging and Rx effect comparisons, also see **Methods**). Inset of left panel: symbol for transcript. **d**, Summary bar plot of the proportions of DEGs under different categories (as specified in the inset of **c**). **e** and **f**, Similar to **c** and **d**, but for showing exenatide treatment effects in aged animals with hypothalamic *Glp1r* knockdown. **g**, Scatter plots showing DNA methylation level changes in aging (*x*-axis) vs. exenatide treatment (*y*-axis) in sites covered by the Illumina mouse 285k array (mm285k) across different tissue organs and circulating WBCs. Each dot represents one differentially methylated locus (DML), color-coded by treatment (Rx) effect categories (as shown in the inset of **c**). Inset of left panel: symbol for DNA methylation. **h**, Similar to **g**, but for showing exenatide treatment effects in aged animals with hypothalamic *Glp1r* knockdown. **i**, Scatter plot showing plasma metabolomic changes in aging (*x*-axis) vs. exenatide treatment (*y*-axis). Each dot represents one metabolite. **j**, Similar to **i**, but for showing exenatide treatment effects in aged animals with hypothalamic *Glp1r* knockdown. In **c**, **e**, and **g**–**j**, the lines represent linear fits (with confidence interval (grey) in **i** and **j**). The Spearman correlation coefficients (*R_S_*) and associated *P*-values are also shown. Sample sizes for data in **c**–**j**: *n* = 5 mice for each experimental group for the various tissues, except *n* = 4 aged exenatide-treated mice (src group) for circulating WBCs transcriptomic profiling.

Consistent with the previous findings, we again observed robust age-counteracting transcriptomic effects of exenatide treatment in the hippocampus, frontal cortex, circulating WBCs, heart, and skeletal muscle (**Fig. 4c, d**). Interestingly, these effects appeared to be even stronger in this cohort of mice where treatment was delivered at an older age (i.e., from 18 to 21 m.o.) (**Fig. 4c, d**) compared to the earlier treatment cohort (**Fig. 2a–h**). The counteraction of age-related transcript expression changes induced by exenatide treatment in the hippocampus remained similarly strong after knocking down hypothalamic *Glp1r* (**Fig. 4c–f, Supplementary Fig. 6e**). However, in the frontal cortex, circulating WBCs, heart, and skeletal muscle, the transcriptomic rejuvenation effects of exenatide treatment were weakened or even abolished by the *Glp1r* knockdown (**Fig. 4e, f, Supplementary Fig. 6e**). Intriguingly, exenatide treatment even exacerbated some age-related transcript expression changes in the heart and skeletal muscle of the hypothalamic *Glp1r* knockdown mice (**Fig. 4e, f, Supplementary Fig. 6e**).

We similarly found global counteraction of aging-associated methylation changes by exenatide treatment in this cohort across the hippocampus, frontal cortex, circulating WBCs, heart, and skeletal muscles (**Fig. 4g**). Such an effect was strongly attenuated in animals with hypothalamic *Glp1r* knockdown in the examined tissue organs and circulating WBCs (**Fig. 4h, Supplementary Fig. 6f**). Notably, this occurred even in the hippocampus (**Fig. 4g, Supplementary Fig. 6f**), despite the preservation of age-counteracting transcriptomic impact (**Fig. 4e, f**). Although the modest age-counteracting effect of exenatide treatment on the circulating metabolome (**Fig. 4i**) appeared to be generally consistent with or without hypothalamic *Glp1r* knockdown (**Supplementary Fig. 6g**), the negative correlation between aging- and exenatide treatment-associated effects was diminished in the mice with *Glp1r* knockdown (**Fig. 4j**).

Based on these results, we concluded that hypothalamic GLP-1R is crucial in mediating the molecular rejuvenating effects of GLP-1RA treatment throughout the body. These data support the notion that the beneficial outcomes of GLP-1RA treatment are dependent on the activation of hypothalamic GLP-1R, which in turn contributes to the systemic rejuvenation.

### GLP-1R agonism and mTOR inhibition modulate multi-omic landscapes along shared and distinct axes in aging

Based on the molecular age-counteracting effects of GLP-1RA and the expected effects of mTOR inhibition^1,57^, we speculated that exenatide and rapamycin treatments may converge substantially in their pharmacological effects in aged mice. To test this, we included an additional experimental group of animals treated with rapamycin (8 mg/kg bw/2 days I.P.), from which we obtained identical metabolic and molecular readouts (**Fig. 4a, b, Supplementary Fig. 6a–d**). Consistent with the known effects of mTOR inhibition^57^, rapamycin treatment led to slight trends of decreased food intake (**Supplementary Fig. 6a**) and body weight (**Supplementary Fig. 6b**), significant impairment in glucose tolerance^58^ (**Supplementary Fig. 6c**), and reduction in gonadal fat (**Supplementary Fig. 6d**).

The age-counteracting effects of rapamycin were evident in the transcriptomes and DNA methylomes of the hippocampus, frontal cortex, circulating WBCs, heart, and skeletal muscle (**Fig. 5a–c**), as well as in the circulating metabolome (**Fig. 5d**). Remarkably, the effects of exenatide treatment correlated well with those observed with rapamycin at transcriptomic, methylomic, and circulating metabolomic levels (**Fig. 5e–g**). Their efficacies appeared similarly potent, as reflected by the transcriptomic and methylomic changes in the hippocampus, circulating WBCs, heart, and skeletal muscle (**Fig. 4c, d, g, Fig. 5a–c, e, f**). In the skeletal muscle, exenatide appeared to induce even strong molecular rejuvenation (**Fig. 4c, d, Fig. 5a, b**). Meanwhile, rapamycin exhibited a stronger impact on the frontal cortex transcriptome (**Fig. 4c, d, Fig. 5a, b**) and the circulating metabolome (**Fig. 4i**, **Fig. 5d, g**).

**Figure 5.**
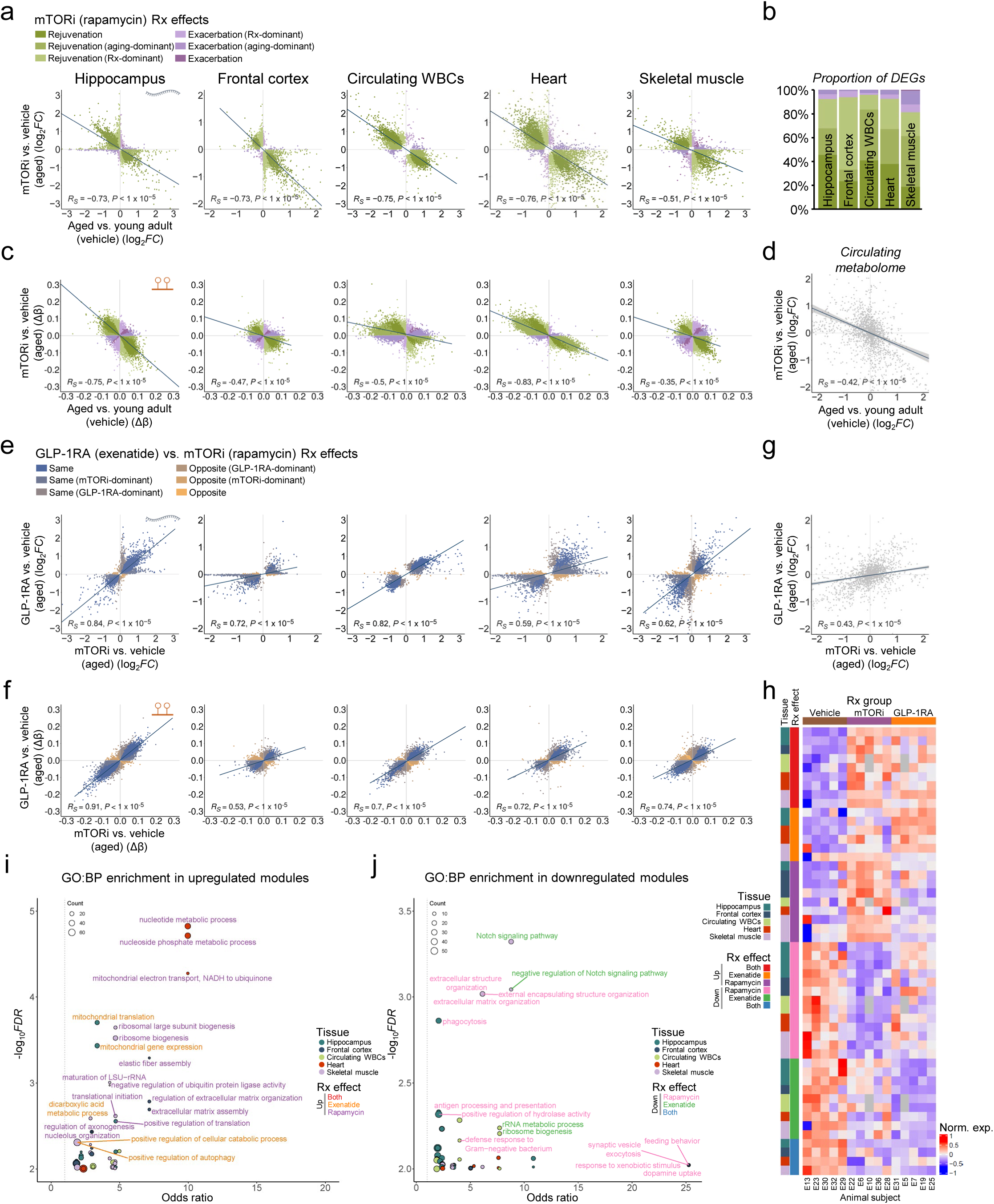
Age-counteracting molecular effects of GLP-1RA (exenatide) treatment vs. that of mTOR inhibition (rapamycin). **a**, Scatter plots showing transcriptomic changes in aging (*x*-axis) vs. rapamycin treatment (*y*-axis) in different tissue organs and circulating white blood cells (WBCs). Each dot represents one differentially expressed gene (DEG), color-coded by treatment (Rx) effect categories as shown in the inset (rejuvenation: opposing aging and Rx effects; exacerbation: same aging and Rx effects; aging- or Rx-dominant: statistically significant in the respective comparison only; no additional label: significant for both aging and Rx effect comparisons, also see **Methods**). Inset of left panel: symbol for transcript. **b**, Summary bar plot of the proportions of differentially expressed genes (DEGs) under different categories (as specified in the inset of **a**). **c**, Scatter plots showing DNA methylation level changes in aging (*x*-axis) vs. exenatide treatment (*y*-axis) in sites covered by the Illumina mouse 285k array (mm285k) across different tissue organs and circulating WBCs. Each dot represents one differentially methylated locus (DML), color-coded by treatment Rx effect categories (as shown in the inset of **a**). Inset of left panel: symbol for DNA methylation. **d**, Scatter plot showing plasma metabolomic changes in aging (*x*-axis) vs. rapamycin treatment (*y*-axis). Each dot represents one metabolite. **e**, **f**, and **g**, Scatter plots showing transcriptomic (**e**), DNA methylation level (**f**), and plasma metabolomic (**g**) changes with rapamycin (*x*-axis) vs. exenatide (*y*-axis) treatment. Inset of **e**, DEG and DML categories, indicating: same or opposite Rx effects; GLP-1RA- or mTORi-dominant: statistically significant in the respective comparison only; no additional label: significant for both Rx effect comparisons (also see **Methods**). Inset of left panels of **e** and **f**: symbols for transcript and DNA methylation, respectively. In **a**, **c**–**g**, the lines represent linear fits (with confidence interval (grey) in **d** and **g**). The Spearman correlation coefficients (*R_S_*) and associated *P*-values are also shown. **h**, Heatmap showing the normalized expression (Norm. exp.) levels of significant differentially expressed gene modules with exenatide and/or rapamycin treatment. Inset (left) shows the color coding for tissue origin and Rx effect category (i.e., significant with exenatide, rapamycin, or both treatments). **i**, Bubble chart showing the gene ontology–biological process (GO:BP) pathways with significant enrichment among the different treatment-upregulated gene modules with exenatide and/or rapamycin treatment (top 3 for each tissue organ/circulating WBCs plotted). Gene count is represented with bubble size. Tissue origin and Rx effect category are color-coded as per the inset. **j**, Similar to **i**, but for treatment-downregulated gene modules. Sample sizes for all plots: *n* = 5 mice for each experimental group for the various tissues, except *n* = 4 aged exenatide- and 4 rapamycin-treated mice for circulating WBCs transcriptomic profiling.

Finally, we performed WGCNA on the transcriptomic data to explore the functional gene modules affected either commonly or distinctively by treatments with exenatide and rapamycin. The analysis corroborated the substantial transcriptomic concordance induced by the two treatments, with numerous shared differentially expressed gene modules (**Fig. 5h**). While we also noted unique gene modules were significantly altered by each treatment, the overall directionality of expression changes was consistent (**Fig. 5h**). Pathway analysis revealed that exenatide preferentially upregulated genes involved in mitochondrial energy metabolism within the hippocampus (**Fig. 5i**) and downregulated Notch signaling-related genes in skeletal muscle (**Fig. 5j**). Conversely, rapamycin more strongly enhanced gene expressions influencing energy metabolism in the heart (**Fig. 5i**). Hence, while exenatide and rapamycin phenocopy each other’s effects, they also modulate age-related molecular changes in a tissue-specific and treatment-dependent manner.

## Discussion

Here, we show that GLP-1R agonism is a promising strategy for the mitigation of age-related decline and extension of healthspan. Encouragingly, the age-counteracting effects of GLP-1RA treatment are widespread, spanning across tissues of the whole body, the circulatory system, and multiple molecular layers as demonstrated by our multi-omic assays. It has been proposed that differentiating between rate, baseline, and combined models in anti-aging interventions allows us to identify whether they target aging mechanisms directly, mimic the targeting of aging processes, or have a mixed effect^59,60^, respectively. Based on our data, the specific functional and molecular responses to exenatide treatment in the aged mice suggest that the rejuvenation effects of GLP-1R agonism align with a rate model (i.e., slowing of aging). Crucially, in our experiments, we selected the dosage of exenatide to minimize potential confounders, in particular aiming to achieve the rejuvenation effects without significantly influencing food intake or body weight. The full spectrum of functional benefits and molecular impacts of GLP-1RA in aging could be even more substantial at higher but still safe and tolerable dosages.

The mechanisms underlying the observed effects are undoubtedly complex, given that systemically administered exenatide activates GLP-1R at various body sites. For example, GLP-1RA enhances the firing of GLP-1R-expressing pro-opiomelanocortin (POMC^GLP-1R+^) neurons in the arcuate nucleus of the hypothalamus^61,62^. The significant reliance of the observed age-counteracting molecular effects on hypothalamic GLP-1R suggests that GLP-1RA-driven rejuvenation may depend on modulating hypothalamic POMC^GLP-1R+^ neuronal activities. In addition, potential systemic impacts mediated by glial GLP-1R activation need to be considered^63,64^. Our findings are in line with previous studies indicating that CNS GLP-1R is an essential component of the signaling network mediating not only neurological^65–67^ but also systemic anti-inflammatory actions of GLP-1RA^68^. The brain, like other body organs, is inevitably impacted by peripheral organ and circulatory changes in aging^69–72^. Conversely, brain aging affects its coordinating role in body-wide processes^73,74^. These all underscore the profound influence of the CNS on systemic health, and suggest that optimizing CNS exposure could be advantageous for the design of new drugs targeting or co-targeting GLP-1R.

Interestingly, in the animals treated from middle age (i.e., 11 m.o. for 30 weeks), we found a divergence in omic impacts in some organs, with the colon DNA methylome being less responsive to GLP-1RA treatment compared to substantial transcriptional changes, and the opposite pattern in the liver and kidneys. Moreover, after hypothalamic *Glp1r* knockdown in even older animals, robust age-counteracting effects in the hippocampus persisted in the transcriptome, but not in the DNA methylation profile. These findings imply that the downstream effects of GLP-1R agonism on transcript expression may involve other epigenetic or gene expression regulatory mechanisms that are body-site-specific. As we observed generally stronger molecular age-counteracting transcriptomic and methylomic effects in the brain, heart, skeletal muscle, and circulating WBCs in the older animals, it remains unclear if the divergence of omic responses to GLP-1R agonism in some organs persists in older age.

We found substantial convergence of multi-omic impacts between exenatide and rapamycin. This raises the need to systematically explore the relative advantages and disadvantages of adopting GLP-1R agonism vs. mTOR inhibition. At the preclinical level, it would be necessary to investigate how each approach might yield different anti-aging efficacies across various organ systems and cell types. Additionally, examining potential overlaps in their mechanisms of action, such as effects on POMC neurons and other modulatory effects in the CNS^75,76^, would be valuable. Past and ongoing clinical studies are actively evaluating the anti-aging potential of mTOR inhibition^77,78^. It remains to be seen whether there are overlapping clinical benefits, taking into account the side effect profiles of each drug class. Although concerns have been raised about skeletal muscle loss with GLP-1R agonists, the strong molecular age-counteracting effects observed with exenatide even when compared to rapamycin in our data suggest the potential to achieve muscle benefits using a low-dose regimen without substantial overall body weight changes.

Given the extensive range of rejuvenation effects observed, it will be important to evaluate the potential of GLP-1RAs in treating human age-related diseases beyond the current clinical indications for diabetes, obesity and their associated comorbidities. Ongoing phase III trials are investigating the use of GLP-1RA in treating Alzheimer’s disease^79^, while we are also conducting a pilot trial focused on cerebral small vessel disease (ClinicalTrials.gov identifier: NCT05356104). Our findings also open the avenue for a nuanced therapeutic strategy that combines a GLP-1RA or mTOR inhibitor with molecular entities that engage disease-specific molecular targets or address issues of selective vulnerability^80^. For instance, combining a GLP-1RA with an anti-amyloid agent^81,82^ for Alzheimer’s disease treatment could be advantageous, where the GLP-1RA may also serve as a pretreatment or as an adjunct therapy once brain amyloid levels are normalized. However, mechanistic compatibility will need to be experimentally verified for each specific combination of therapeutic approaches and in individual clinical conditions. This approach may allow us to leverage synergistic anti-aging and neuroprotective effects to combat complex, multifactorial age-related conditions.

Collectively, our work has provided multifaceted evidence for a comprehensive body-wide anti-aging strategy based on GLP-1R agonism, which, given its current clinical use and favorable safety profile, introduces an imminently deployable anti-aging intervention. Although we have not demonstrated lifespan extension in mice, the strong similarities with the effects of mTOR inhibition, which is well-proven for its longevity benefits^1,57^, suggest the potential for GLP-1R agonism to enhance lifespan. We also acknowledge that we only performed experiments in male mice, while any anti-aging intervention could in principle exhibit gender dependence. However, given that the clinically reported cardiorenal^83–85^ and pleiotropic^32–34,36^ benefits of GLP-1RAs were similar across sexes, the age-counteracting effects of GLP-1R agonism are likely consistent across different genders. It is also conceivable that GLP-1R agonism may complement other anti-aging or rejuvenation methods. Future longitudinal studies are necessary to explore these possibilities. To further elucidate the cellular mechanisms behind the rejuvenation observed with GLP-1RA treatment, subsequent research could also employ single-cell and spatial genomic profiling, along with appropriate functional and histological assays. These will help to identify cell-type-specific rejuvenation effects, clarify the full impact of GLP-1R agonism on aging, and inform the design of clinical trials.

## Methods

### Animals

All experimental procedures were approved in advance by the Animal Experimentation Ethics Committee of the Chinese University of Hong Kong (CUHK), and were carried out in accordance with the Guide for the Care and Use of Laboratory Animals. All C57BL/6 mice were bred and provided by the Laboratory Animal Service Center of CUHK unless otherwise specified, and maintained at controlled temperature (22–23 °C) with an alternating 12-hour light/dark cycle with free access to standard mouse diet and water. The ambient humidity was maintained at < 70 % relative humidity.

Three cohorts of mice were used. For the aged long-term treatment cohort, mice were treated with the GLP-1RA exendin-4 or vehicle from 11 months old (m.o.) for 30 weeks (see next section for details of drug and vehicle treatments). A group of 3 m.o. mice that received vehicle treatment for 4 weeks were included as a control group. For the young long-term treatment cohort, mice were treated with exendin-4 or vehicle from 3 m.o. for 26 weeks. For the aged short-term treatment cohort, mice were imported from the Jackson Laboratory and received adenovirus-associated virus (AAV) vector-mediated hypothalamic *Glp1r* knockdown or a control AAV injection (see below for details). Mice were allowed to recover from AAV injection for 8 weeks, and subsequently subjected to vehicle, exendin-4 or rapamycin treatment from 18 m.o. for 13 weeks. A control group of young mice underwent a control AAV injection and 4 weeks recovery period, and then received vehicle treatment from 5 m.o. for 7 weeks.

### Exendin-4 and rapamycin treatment

Exendin-4 (HY-13443, MedChem Express, China) was reconstituted in phosphate-buffered saline (PBS) to 1 mg/ml and further diluted in PBS to 4.2 µg/ml. For all GLP-1RA treatment groups, each mouse received daily intraperitoneal (I.P.) injections of exendin-4 at a dosage of 21 µg/kg body weight (bw) (equivalent to 5 nmol/kg bw/day). Rapamycin (HY-10219, MedChem Express, China) was dissolved in absolute ethanol to 10 mg/ml and subsequently diluted in a vehicle solution consisting of 5% Tween 80, 5% PEG-400, and PBS to 0.8 mg/ml. For the mTOR inhibitor treatment group, each mouse received I.P. injections of rapamycin at a dosage of 8 mg/kg bw on alternate days (equivalent to 8.75 µmol/kg bw/2 days). Mice in the vehicle control groups received daily volume-matched I.P. PBS injections.

### AAV-mediated hypothalamic GLP-1R knockdown

Short hairpin RNA (shRNA) (sequence: 5’-GCGTCAACTTTCTTATCTTCA-3’) for *Glp1r* knockdown was constructed into the miR-30 scaffold^86^ driven by the EF1a promoter, and packaged into recombinant AAV-2/9. Control AAV contained a scrambled sequence (5’-CCTAAGGTTAAGTCGCCCTCG-3’). Young (4 m.o.) and aged (16 m.o.) mice were anesthetized by I.P. injection of 150 mg/kg bw ketamine and 10 mg/kg bw xylazine, and positioned in a stereotaxic frame (Model 68528, RWD Life Science, China) for injection. A total of 1 microliter AAV (containing 5–6 × 10^9^ vg) was injected into the hypothalamus of each mouse. To ensure a broad regional coverage of the hypothalamus, AAVs were delivered to four sites, with coordinates as follows: anteroposterior (AP) = -1.6 mm, mediolateral (ML) = ±0.25 mm, dorsoventral (DV) = -5.9 mm from the dura (0.2 µl per site), and AP = -2 mm, ML = ±0.25 mm, DV = -5.85 mm from the dura (0.3 µl per site). To verify *Glp1r* transcript knockdown efficiency with quantitative PCR, brains were removed after PBS perfusion and half of the dissected hypothalamus tissues from 3 mice per experimental group were used for RNA extraction using TRIzol reagent (15596018, Thermo Fisher Scientific, US). Primers used for transcript amplification were: *Glp1r* (forward: 5’-CAGTGGGGTACGCACTTTCT-3’; reverse: 5’-TAACGAACAGCAGCGGAACT-3’); *Gapdh* (forward: 5’-CATCTTCCAGGAGCGAGACC-3’; reverse: 5’-GGCGGAGATGATGACCCTTT-3’).

### Physical and cognitive assessments

A grip strength meter (Model XR501, Shanghai Xin-Ruan Instruments Inc., Shanghai, China) was used to measure forelimb grip strength. The mouse subject was allowed to grip the mesh lattice with its forelimbs. The peak force to pull the mouse away from the grip was recorded. For each mouse, the average value from five repeated measurements was obtained, with 15-minute intervals between successive measurements.

The accelerated rotarod test equipment (Model R03-1, Xin-Ruan Instruments Inc., Shanghai, China) consisted of a 3-cm diameter rod. Mice were trained to habituate to the rotating rod by walking on it at a low constant speed (4 rotations per minute (R.P.M.), 2 min × 3 sessions with 15-min intervals at the baseline, and 2 min × 1 session at the 3- or 6-month treatment time points). During testing, rod rotation was linearly accelerated from 4 to 40 R.P.M over 5 minutes. The endpoint was reached when the mouse could not withstand the rotation and fell from the rod, or resisted falling over 5 minutes. For each mouse, the average duration to reach the endpoint in five trials was obtained, with 15-minute intervals between successive trials.

A home-made Barnes maze was used, which consisted of a white acrylic round disk (90-cm diameter) with 20 holes (5-cm diameter) radially evenly spaced at 5 cm from the outer edge. 19 of the 20 holes were blocked while the remaining one provided access to an escape box. Visual cues were placed around the maze in the testing room. On the first day, mice were first trained to find the escape box. At the beginning of the assay, the mouse subject was covered by a cardboard box at the center of the maze. Five seconds later, an aversive noise was initiated from a nearby device (900 Hz, 80 dB), and the box was removed to permit exploration and escape. Entering the escape box or spending at least five seconds exploring the correct escape hole was defined as an escape event. Mice that failed to find the escape box in 3 minutes were guided to it using a glass cylinder. One hour later, mice were tested by the same procedure, and the so-obtained values were used as the performance measures for the animals for assay day one. On each day from assay days two to four, mice were tested the same way twice with a one-hour interval, and for each animal the two measurements were averaged to give the performance measure.

The open field test was performed in a home-made 56 cm × 56 cm arena with opaque walls. The mouse was placed in the arena and allowed to freely explore for 10 minutes, while its locomotion was recorded from above using a camera (StreamCam, Logitech, China). The total distance traveled was extracted from the video using the open-source package MouseActivity^87^ (https://github.com/HanLab-OSU/MouseActivity) in MATLAB R2022a (MathWorks, US).

To evaluate longitudinal treatment effects on forelimb grip strength and rotarod performance, the corresponding measurements relative to vehicle control group for each mouse were calculated by subtracting the values at the corresponding time points by the corresponding group mean values of vehicle-treated mice. One-sided Page’s trend test was used to examine the presence of any significant increasing trends observed in the treatment group. The performance of animal subjects in the Barnes maze across treatment groups over the four assay days was analyzed using two-way repeated-measures ANOVA. Comparison between the locomotor activity levels of the vehicle- and exendin-4-treated groups was conducted using two-sided, unpaired *t*-test.

### Metabolic assessment

The body weight of each mouse was monitored on a weekly basis. The mice in each treatment group were housed together in a single cage. The food provided to each cage was weighed once weekly and averaged to determine the daily food intake per mouse. For the long-term (30-week) treatment experiment, the monitoring started at baseline (one week before starting treatment) and continued until the conclusion of the study. For the short-term (13-week) treatment experiment, it began at baseline and was conducted for seven weeks following the initiation of treatment. The impact of treatments on longitudinal body weight was assessed using two-way repeated-measures ANOVA.

Drug or vehicle administration was halted on the day of fasting blood glucose measurement or oral glucose tolerance test (OGTT). Mice underwent a 6-hour fasting period prior to fasting blood glucose measurement or OGTT. For OGTT, blood glucose levels were measured from tail vein samples at the baseline, and then at 15, 30, 60, and 120 minutes after administration of 2 g/kg bw of glucose via oral gavage. Comparison of fasting blood glucose levels in the vehicle- vs. exendin-4-treated groups, and at baseline vs. after 6-month treatment, was conducted using two-way repeated measures ANOVA. Area under the curve (AUC) values were calculated from the measurements obtained from OGTT, and compared across experimental groups using one-way ANOVA with Holm-Sidak’s post-hoc multiple comparisons test.

### Tissue collection and storage

On the day of tissue collection, mice received the final dose of vehicle or drug treatment in the morning, followed by a 6-hour fasting period. Mice were euthanized via isoflurane inhalation and transcardially perfused with 20 ml of ice-cold PBS. Mouse brains were removed from the cranium and underwent further manual microdissection to isolate the hypothalamus, hippocampus, and frontal cortex bilaterally. From the heart, the left ventricle was collected for RNA extraction, and the left atrium for DNA extraction. The quadriceps femoris muscle was collected as the skeletal muscle sample. For the colon, the proximal segment was collected. The whole liver, spleen, bilateral kidneys and lungs were dissected and parts of the tissue were collected. Bilateral gonadal adipose tissues were collected, and weighed prior to further processing (aged long-term treatment cohort: *n* = 3 and 6 mice had the adipose tissues weighted for vehicle control and exenatide-treated groups respectively; for all other cohorts, adipose tissue weighting was done for all animal subjects). The collected tissues from different organs were further dissected into smaller pieces, each with largest dimension <5 mm and preserved in RNA*later* (AM7021, Thermo Fisher Scientific, US) for RNA extraction, or RNA/DNA shield (R1200-125, Zymo Research, US) for DNA extraction. All samples were stored at -20°C before extraction.

The blood samples were lysed in an ice-cold red blood cell lysis buffer for 20 minutes. The white blood cells were separated via centrifugation at 500 ×g, 4°C for 10 minutes, and subsequently lysed into TRIzol reagent (15596018, Thermo Fisher Scientific, US) for future RNA extraction, or genomic DNA lysis buffer (BioFluid & Cell Buffer, Zymo Research, US) for DNA extraction. Plasma was isolated from EDTA-anticoagulated blood samples by centrifugation at 1,600 ×g, 4°C for 15 minutes. The samples were stored at -80°C before RNA/DNA extraction or further assays.

### Total RNA and genomic DNA extraction

The tissues were retrieved from RNA*later* (AM7021, Thermo Fisher Scientific, US), combined with the TRIzol reagent (15596018, Thermo Fisher Scientific, US) or the lysis buffer of the RNAqueous kit (AM1912, Thermo Fisher Scientific, US), homogenized using the TissueLyser II system (Qiagen, US), and total RNA extraction was performed following the instructions in the vendor’s manuals. Genomic DNA was extracted using a similar procedure, with blood DNA extracted with the Quick-DNA Miniprep Plus Kit (D4069, Zymo Research, US) and solid tissue DNA extracted using the E.Z.N.A. Tissue DNA Kit (D3396-02, Omega Bio-Tek, US).

### Plasma metabolomic measurement

Plasma metabolomic profiling was performed using high-throughput liquid chromatography-mass spectrometry (LC-MS) at BGI Genomics Inc. (Shenzhen, China). To extract metabolites, 100 µl of plasma from each mouse was combined with 700 µl of extractant containing internal standard (methanol:acetonitrile:water = 4:2:1, v/v/v), shaken for 1 minutes and kept at -20℃ for 2 hours. The samples were then centrifuged at 25,000 ×g, 4℃ for 15 minutes. The supernatant was collected and the solvent was dried out. The pellet was subsequently reconstituted in 180 µl of methanol:water (1:1 v/v) and the sample was centrifuged again at 25,000 ×g, 4℃ for 15 minutes. The resulting supernatant was subjected to LC-MS/MS analysis, using the Waters UPLC I-Class Plus equipped with a Waters ACQUITY UPLC BEH C18 column (1.7 µm, 2.1 mm × 100 mm) (Waters, US) and tandem Q-Exactive high resolution mass spectrometer (Thermo Fisher Scientific, US). Pooled plasma reference samples, which were prepared by combining small aliquots from the study samples, were analyzed among the participant samples to monitor the repeatability of the analysis process.

The column temperature was maintained at 45℃. The mobile phase consisted of 0.1% formic acid (A) and acetonitrile (B) in the positive mode, and in the negative mode, the mobile phase consisted of 10 mM ammonium formate (A) and acetonitrile (B). The gradient conditions were as follows: 0–1 min, 2% B; 1–9 minutes, 2%–98% B; 9–12 minutes, 98% B; 12–12.1 minutes, 98% B to 2% B; and 12.1–15 minutes, 2% B. The flow rate was 0.35 ml/min and the injection volume was 5 μl. The full scan range was 70–1050 m/z with a resolution of 70000, and the automatic gain control (AGC) target for MS acquisitions was set to 3 × 10^6^ with a maximum ion injection time of 100 ms. Top 3 precursors were selected for subsequent MS-MS fragmentation with a maximum ion injection time of 50 ms and resolution of 17500, with the AGC set at 1 × 10^5^. The stepped normalized collision energy was set to 20, 40 and 60 eV. ESI parameter settings were: sheath gas flow rate 40, aux gas flow rate 10, positive-ion mode spray voltage (|KV|) 3.80, negative-ion mode spray voltage (|KV|) 3.20, capillary temperature 320°C, aux gas heater temperature 350°C.

### Bioinformatic data analysis

All analysis was performed with R (v4.2.2.) unless otherwise specified.

### Metabolome data analysis

The mass spectrometry data were processed by BGI Genomics Inc. (Shenzhen, China) using the Compound DiscoverTM 3.3 software (Thermo Fisher Scientific, US) and analyzed in combination with the BGI metabolome, mzCloud (https://www.mzcloud.org/), and ChemSpider^88^ online databases. A data matrix containing information, including metabolite identification and peak area was obtained for further analysis.

The normalized intensities further underwent variance-stabilizing transformation (VST) to account for systematic bias^89,90^. After mean variance stabilization, differential analysis was performed using linear models as implemented within the limma package (v3.54.2) in R. Spearman correlation coefficients and *P*-values were calculated by correlating the log fold changes per metabolite in aged drug-treated vs. aged vehicle-treated mice and in aged vehicle-treated vs. young vehicle-treated mice.

### RNA-seq data analysis

Raw sequencing FASTQ files were mapped with STAR (v2.7.10b) with default parameters using the *Mus musculus* genome assembly GRCm39. Quality control was carried out with the FastQC (v.0.11.9) and MultiQC (v.1.13a) packages. HTSeq (v.2.0.2) was used to obtain read counts with default options except ‘mode’ being set to ‘intersection-strict’, and ‘nonunique’ being set to ‘fraction’ for ambiguous reads. Gene counts were filtered for a minimum of at least 10 reads in a minimum number of samples, with the minimum number based on the number of animals constituting the smallest comparator group size in the experiment.

Downstream analysis was done with custom scripts run on R (v.4.2.2 and v4.3.1). Read counts were normalized with VST for exploratory data analysis and heatmap visualizations. Differential expression analysis was performed with DESeq2 (v.1.38.3). Log fold change estimates were shrunken with the approximate posterior estimate for generalized linear models method. Significant differentially expressed genes (DEGs) were taken as those with a false discovery rate (FDR)-adjusted *P*-value < 0.05. We grouped the DEGs into six categories according to their pattern of differential expression by comparing DEGs in five combinations of contrasts, namely 1) aged GLP-1RA-treated vs. aged vehicle-treated mice against aged vehicle-treated vs. young vehicle-treated mice, 2) the same comparison but with hypothalamic *Glp1r* knockdown, 3) aged GLP-1RA-treated vs. aged vehicle-treated mice against aged mTORi-treated vs. aged vehicle-treated mice, 4) young adult GLP-1RA-treated vs. young adult vehicle-treated mice against aged GLP-1RA-treated vs. aged vehicle-treated mice, and 5) aged GLP-1RA-treated vs. aged vehicle-treated mice (with hypothalamic *Glp1r* KD) against aged GLP-1RA-treated vs. aged vehicle-treated mice (with hypothalamic scramble shRNA AAV injection).

For the first two contrasts, the DEGs were labeled “rejuvenation” where the GLP1-RA treatment-associated change and the age-associated change were in the opposite direction or “exacerbation” where the GLP-1RA treatment-associated change was in the same direction as the age-associated change. An additional label of “Rx-dominant” was given for where only the GLP-1RA treatment associated change reached statistical significance, and of “aging-dominant” was given for where only the age-associated change reached statistical change. No additional label was given if both comparisons were significant. For the third comparison, the patterns were labeled “same” where the GLP-1RA treatment-associated change and mTORi treatment-associated change were in the same direction, and “opposite” when the changes were in the opposite direction. We further specified the dominant agent depending on which treatment yielded the statistically significant change. No additional label was given if both comparisons were significant. For the fourth comparison, the patterns were labeled “same” where the GLP-1RA treatment-associated changes in young adult and in aged mice were in the same direction, and “opposite” when the changes were in the opposite direction. We further specified the dominant effect depending on which age group treatment yielded the statistically significant change. No additional label was given if both comparisons were significant. For the fifth comparison, the patterns were labeled “same” where the GLP-1RA treatment-associated changes in aged mice with hypothalamic *Glp1r* KD and scramble shRNA AAV injection were in the same direction, and “opposite” when the changes were in the opposite direction. We further specified the dominant effect group depending on which comparison yielded the statistically significant change. No additional label was given if both were significant. Spearman correlation coefficients and *P*-values for each contrast were calculated by correlating the log fold changes per gene in the pair of comparisons.

### Weighted gene co-expression network analysis

WGCNA (v.1.72-5) was conducted with the DESeq2 vst-normalized count matrix. For each tissue type, a signed-hybrid network was constructed with biweight midcorrelation, with the soft threshold chosen dynamically for each tissue at the level at which the scale-free topology model fit reached 0.8. The minimum module size was set at 30. Module eigengenes (i.e., module first principal components) were used to summarize modules, and modules with eigengene correlation < 0.3 were merged.

Differential eigengene analysis was performed by comparing the eigengene expressions of each sample across comparator groups with a Wilcoxon rank-sum test or a Kruskal-Wallis test. An FDR of < 0.1 was taken to denote a significant difference in module eigengene expression across treatment groups.

### Pathway enrichment analysis

Pathway enrichment was done with ClusterProfiler (v4.6.2) against the Gene Ontology^91^ and KEGG^92^ databases, and custom gene sets of functional pathways related to aging^93–99^. For each tissue, all genes that underwent differential expression analysis were used as the background gene set.

In pathway enrichment analyses for DEGs, “rejuvenation” DEGs (significant for at least one comparison) were used (ClusterProfiler enrichGO). To assist in pathway enrichment analysis interpretability, enriched terms from the comparisons were visualized using the simplifyEnrichment package (simplifyGOFromMultipleLists).

For gene modules from WGCNA, the genes with a correlation of > 0.7 with the module eigengene were used for the foreground in custom enrichment analysis (ClusterProfiler enricher).

### DNA methylation data analysis

DNA methylation assays were carried out by the Clock Foundation using a custom BeadChip array containing loci from the Infinium Mouse Methylation BeadChip (i.e., mm285k array^56^) and the mammalian methylation array (i.e., mm40k array^54^). Data analysis was performed with Sesame (v1.16.1). In brief, raw IDAT files were read with the Sesame prepSesameData function, with the SHCDPB masks (inferSpecies, prefixMaskButC, inferInfiniumIChannel, dyeBiasNL, pOOBAH, noob). This retained 279,644 probes from the mm285k array for downstream analysis. Differential methylation testing per locus was performed with the normalized β values using mixed linear models as implemented in the Sesame package via the DML function. To analyze methylation sites based on the mm40k array, we obtained an imputed β-value data matrix provided by the Clock Foundation, containing 36,793 probes. We then similarly ran the differential analysis.

The differentially methylated loci (DML, nominal *P*-value < 0.05) were categorized based on the differential pattern across various combinations of contrasts, namely 1) aged GLP-1RA-treated vs. aged vehicle-treated mice against aged vehicle-treated vs. young vehicle-treated mice, 2) the same comparison but with hypothalamic *Glp1r* knockdown, 3) aged GLP-1RA-treated vs. aged vehicle-treated mice against aged mTORi-treated vs. aged vehicle-treated mice, and 4) aged GLP-1RA-treated vs. aged vehicle-treated mice (with hypothalamic *Glp1r* KD) against aged GLP-1RA-treated vs. aged vehicle-treated mice (with hypothalamic scramble shRNA AAV injection).

For the first two contrasts, the DMLs were labeled “rejuvenation” where the GLP1-RA treatment-associated change and the age-associated change were in the opposite direction, or “exacerbation” where the GLP-1RA treatment-associated change was in the same direction as the age-associated change. An additional label of “Rx-dominant” was given for where only the GLP-1RA treatment associated change reached statistical significance, and of “aging-dominant” was given for where only the age-associated change reached statistical change. No additional label was given if both comparisons were significant. For the third comparison, the patterns were labeled “same” where the GLP-1RA treatment-associated change and mTORi treatment-associated change were in the same direction, and “opposite” when the changes were in the opposite direction. We further specified the dominant agent depending on which treatment yielded the statistically significant change. No additional label was given if both comparisons were significant. For the fourth comparison, the patterns were labeled “same” where the GLP-1RA treatment-associated changes in aged mice with hypothalamic *Glp1r* KD and scramble shRNA AAV injection were in the same direction, and “opposite” when the changes were in the opposite direction. We further specified the dominant effect group depending on which comparison yielded the statistically significant change. No additional label was given if both were significant. Spearman correlation coefficients and *P*-values for each contrast were calculated by correlating the delta β value (Δβ) per locus in the pair of comparisons.

## Acknowledgements

We thank Prof. Tom Mrsic-Flogel and Prof. Tom Otis for helpful discussions about the study. We thank Dr. Phil Xie for advice on DNA methylation analysis and helpful comments on the manuscript. We thank Anki Miu, Dorothy Ieong, Becky Yung, and Florence Yau for administrative support to the project. We acknowledge access to TissueLyser II at Prof. Jun Yu’s lab, behavioral box at Dr. Hei Ming Lai’s lab, and support for use of QuantStudio 12K Flex by the core facility management team of the Li Ka Shing Institute of Health Sciences. This work was supported by a Croucher Innovation Award (CIA20CU01) from the Croucher Foundation; the General Research Fund (14100122), the Collaborative Research Fund (C6027-19GF, C7074-21GF and C4062-22EF), and the Area of Excellence Scheme (AoE/M-604/16) of the Research Grants Council, the University Grants Committee of Hong Kong; the Excellent Young Scientists Fund (Hong Kong and Macau) (82122001) from the National Natural Science Foundation of China; the Asian Young Scientist Fellowship from the Future Science Awards Foundation; the Lo’s Family Charity Fund Limited.

## Author Contributions

J.H. and H.K. conceived the study, and designed the research with A.J.K. and V.C.T.M.. J.H. led and conducted all experiments and analyzed the data on behavioral and metabolic readouts. A.J.K. led and conducted bioinformatics data analysis, with input from J.H., and N.L.. J.L. contributed to tissue collection, DNA and RNA extraction, and analysis of transcriptomic data. C.C. contributed to in vivo treatment, behavioral assays, and associated data analysis, and tissue collection. A.T., D.Z., K.C., and M.L. contributed to tissue and blood sample processing. R.C.H.C., Z.W., and M.W. contributed to codes for analyzing video recordings of behavioral assays. L.Y.C.Y. and D.C.W.C. assisted with experimental procedures. B.Y.I., Z.L., W.K.K.W., K.H.-M.C., W.-J.L., Y.T., B.W.-L.N., S.H.W., and T.W.L. contributed to data interpretation and technical discussions. J.H., V.C.T.M., and H.K. acquired funding, supervised and managed the project. J.H., A.J.K., J.L., and H.K. prepared figures. J.H., A.J.K., V.C.T.M., and H.K. wrote the paper with input from all authors.

## Data Availability

Pre-processed data will be made available upon the acceptance of the manuscript.

## Code Availability

Custom-written codes for data analysis will be made available upon the acceptance of the manuscript.

---

**Supplementary Figure 1.**
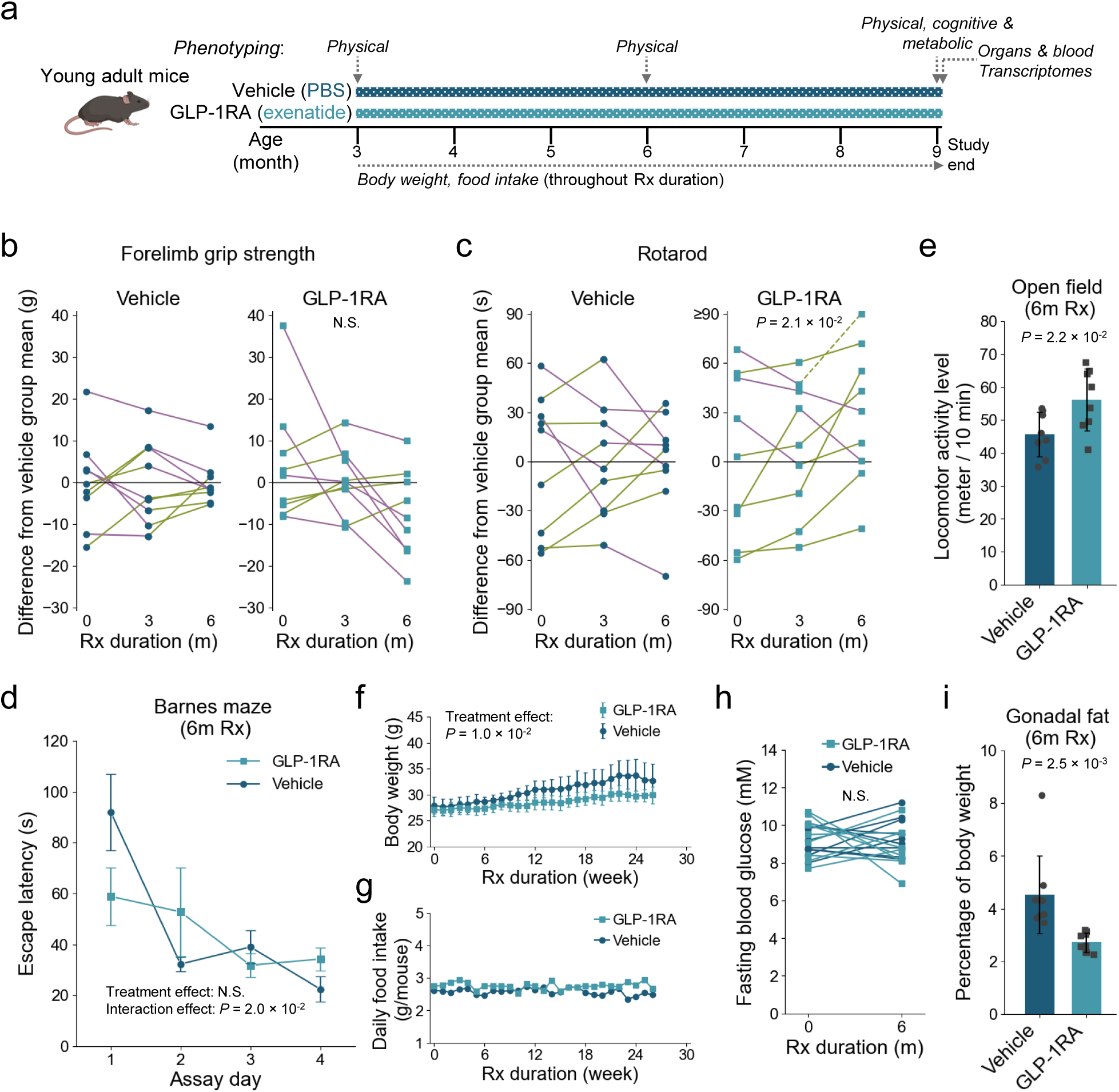
Physical, cognitive and metabolic readouts in young adult mice treated with a GLP-1RA (exenatide) for 30 weeks starting at 3 months old. **a**, Schematic of experimental design for the young adult long-term treatment cohort. **b**, The relative performances of same-age mice treated with exenatide vs. phosphate-buffered solution (PBS) vehicle for the same durations in forelimb grip strength test. Values are plotted as differences from vehicle group mean at each treatment duration time point. Dots connected by lines are longitudinal data from individual animal subjects. The lines are color-coded according to temporal change across two consecutive time points of assessment (green: increase; magenta: decrease). **c**, Similar to **b**, but for performance in the accelerated rotarod test, reflecting motor coordination ability. **d**, Comparison of the performances of exenatide and vehicle-treated mice in the Barnes maze test, a spatial memory-dependent task, after 6-month treatments. Mean (±S.E.M.) time to escape for animals over four consecutive assay days are shown for each treatment group. Shorter times indicate better performances. **e**. Comparison of locomotor activity during a 10-minute open field test after 6-month treatments. **f**, Mean (±S.D.) body weight of the two animal groups throughout treatment period (monitored weekly). **g**, Average daily food intake per mouse for the two animal groups throughout treatment period (monitored weekly). **h**, Fasting blood glucose levels at baseline and after 6-month treatments for each animal subject in the two groups. **i**, Gonadal fat weight as percentage of body weight in the two treatment groups. Sample sizes: *n* = 9 mice for each experimental group in all plots. *P*-values: **b** and **c**, Page’s trend test; **d**, two-way repeated measures ANOVA; **e**, two-sided unpaired *t*-test; **f**, and **h**, two-way repeated measures ANOVA; **i**, two-sided unpaired *t*-test. N.S. in **b**, **d**, and **h** indicate statistical non-significance with *P-*value > 0.05. Abbreviation: Rx, treatment.

**Supplementary Figure 2.**
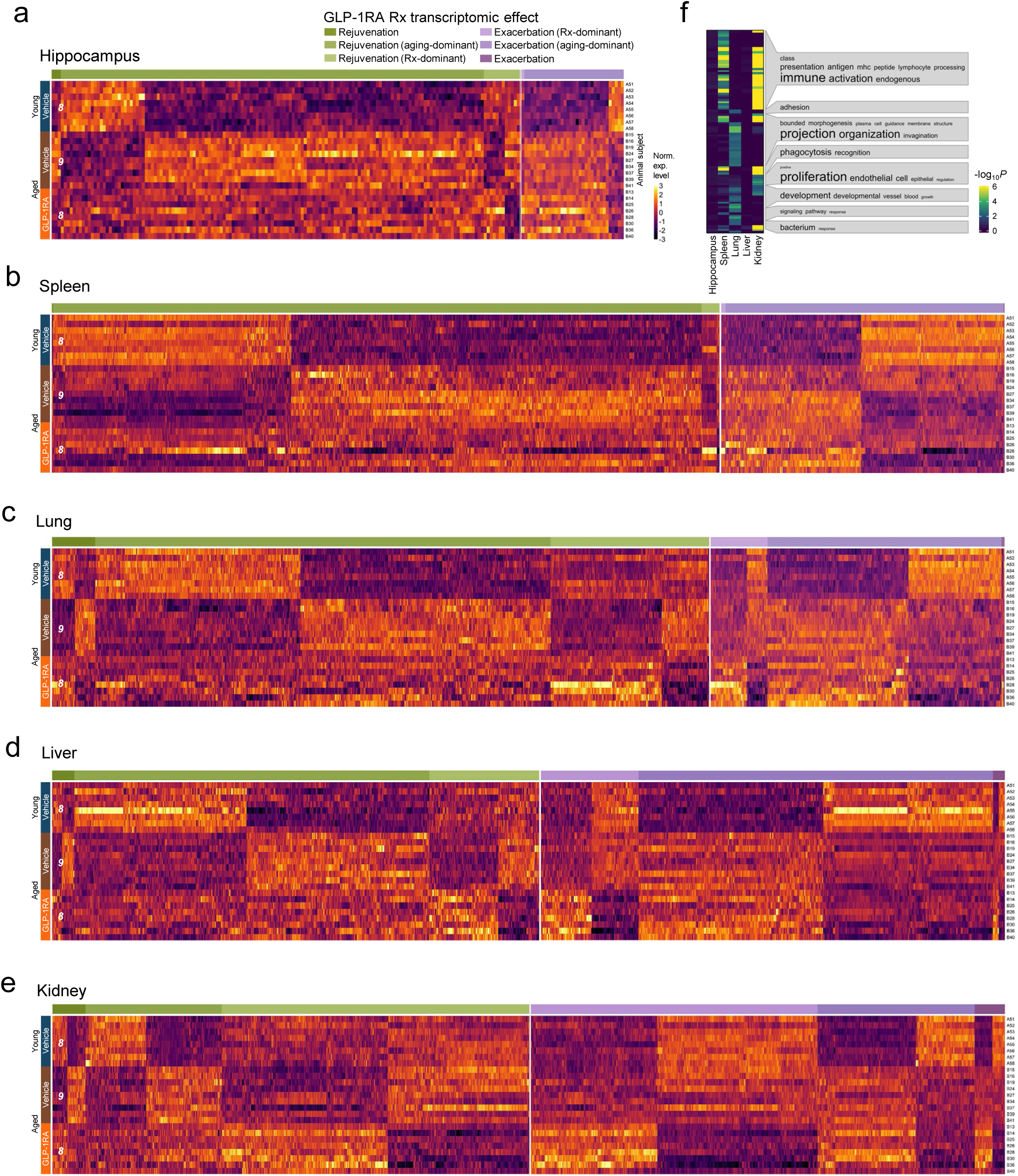
Transcriptomic impacts across other tissues and organs in aging mice treated with a GLP-1RA (exenatide) for 30 weeks starting at 11 months old. **a**–**e**, Heatmaps of the expression levels of protein-coding genes with significant aging-associated and/or exenatide treatment-induced differential expressions in the different tissue organs. The genes are grouped according to the different treatment (Rx) effect categories as specified in the inset of **a** (rejuvenation: opposing aging and Rx effects; exacerbation: same aging and Rx effects; aging- or Rx-dominant: statistically significant in the respective comparison only; no additional label: significant for both aging and Rx effect comparisons, also see **Methods**). Number at the left of each heatmap: samples sizes (numbers of mice) for each tissue. **f,** Functional pathway terms enrichment among aging-associated transcripts with differential expressions that were antagonized by exenatide treatment across the different tissue organs.

**Supplementary Figure 3.**
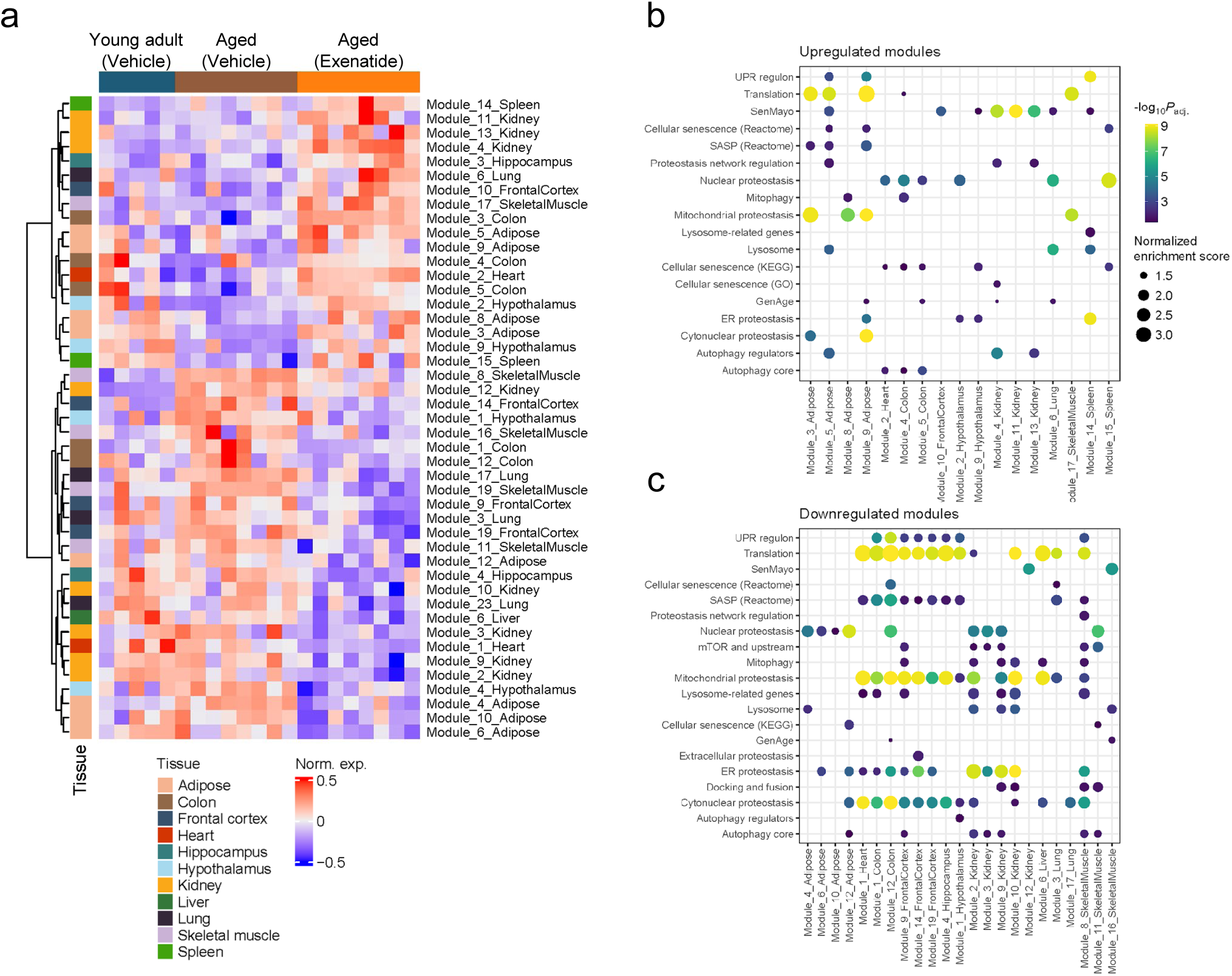
Weighted gene co-expression network analysis (WGCNA) and module gene enrichment analysis for 30-week GLP-1RA (exenatide) treatment effects in aging mice. **a**, Heatmap of the normalized expression (Norm. exp.) levels of significant differentially expressed gene modules with exenatide treatment across animals in the different experimental groups (*n* = 5, 8, and 8 mice for the young adult vehicle, aged vehicle, and aged exenatide treatment groups, respectively, for this analysis). **b** and **c**, Dot plots showing the enrichment of gene sets associated with various hallmarks of aging, among the exenatide treatment-induced differentially expressed modules in the tissue organs.

**Supplementary Figure 4.**
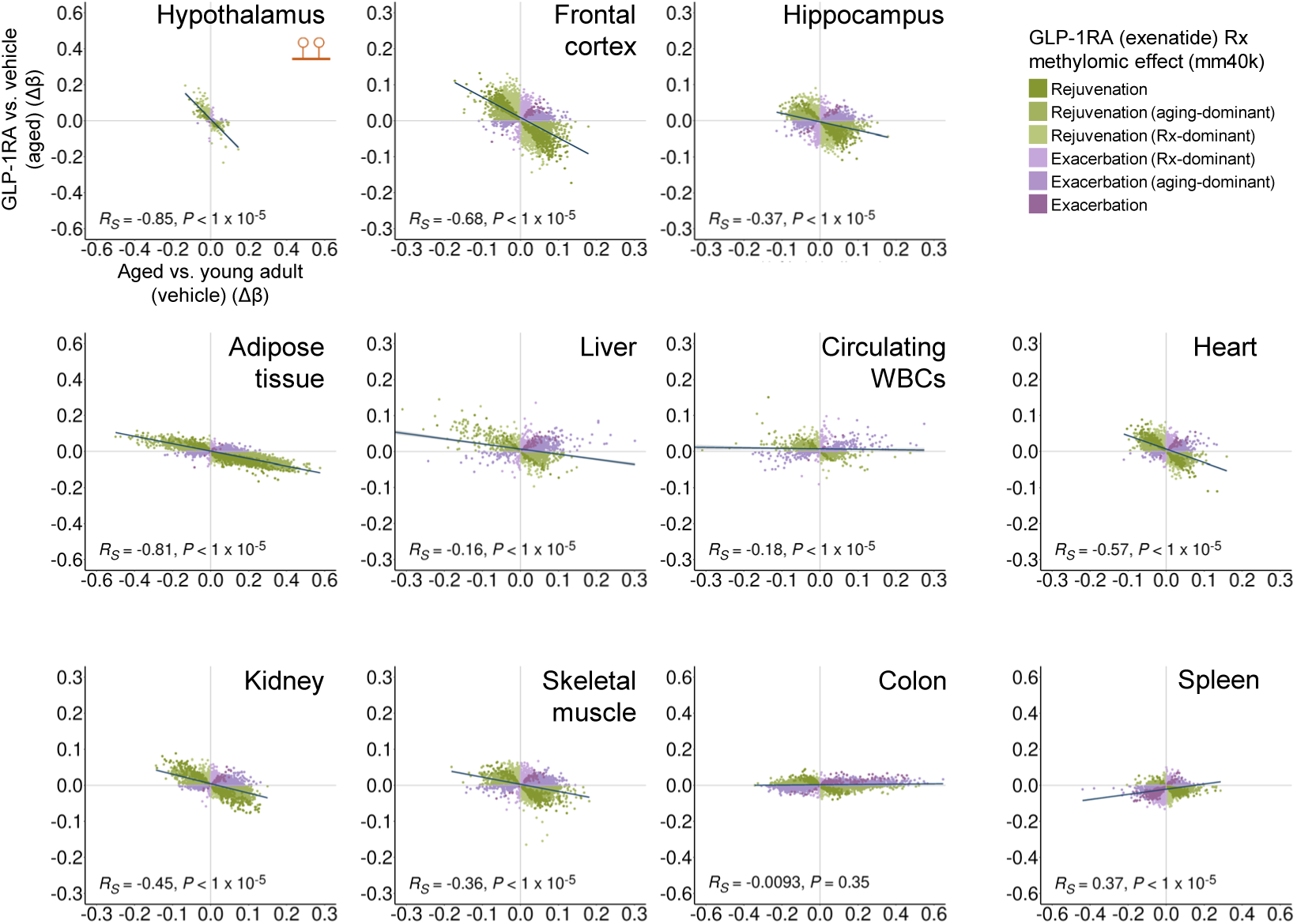
Long-term (30-week) GLP-1RA (exenatide) treatment-induced methylomic changes in aging mice, with analysis restricted to ∼40 thousand mammalian conserved sites. Scatter plots showing DNA methylation level changes in aging (*x*-axis) vs. exenatide treatment (*y*-axis) in imputed sites covered by the mammalian 40k array (mm40k) across different tissue organs and circulating white blood cells (WBCs). Each dot represents one differentially methylated locus (DML), color-coded by treatment (Rx) effect categories as shown in the inset (rejuvenation: opposing aging and Rx effects; exacerbation: same aging and Rx effects; aging- or Rx-dominant: statistically significant in the respective comparison only; no additional label: significant for both aging and Rx effect comparisons, also see **Methods**). For each plot, the line represents linear fit. The Spearman correlation coefficients (*R_S_*) and associated *P*-values are also shown. Inset of left upper panel: symbol for DNA methylation. Sample sizes for the various tissue organs are the same as that of Fig. 3.

**Supplementary Figure 5.**
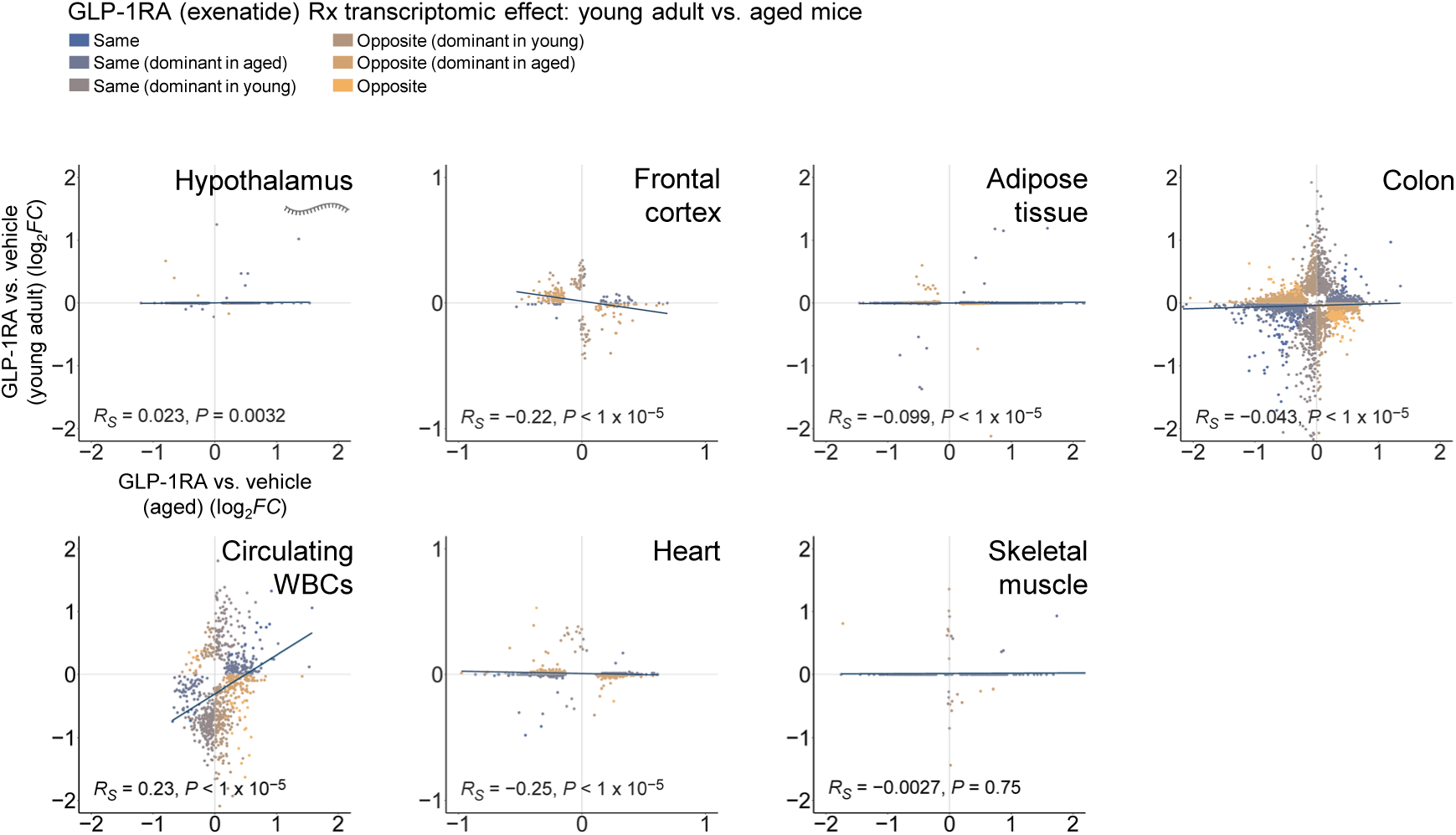
Correlation of transcriptomic impacts across tissues and organs in young adult vs. aged mice treated with a GLP-1RA (exenatide) for 30 weeks. Scatter plots showing transcriptomic changes with exenatide treatment in aged (*x*-axis) vs. young adult (*y*-axis) mice across the different tissue organs and circulating white blood cells (WBCs). Each dot represents one differentially expressed gene (DEG), color-coded by treatment (Rx) effect categories as shown in the inset (same or opposite Rx effects; dominant in aged or young: statistically significant in the respective comparison only; no additional label: significant for both comparisons, also see **Methods**). For each plot, the line represents linear fit. The Spearman correlation coefficients (*R_S_*) and associated *P*-values are also shown. Sample sizes for the aged long-term treatment groups are the same as that in Fig. 2. Inset of left upper panel: symbol for transcript. Sample sizes for the young adult long-term treatment groups: *n* = 8 mice for each group.

**Supplementary Figure 6.**
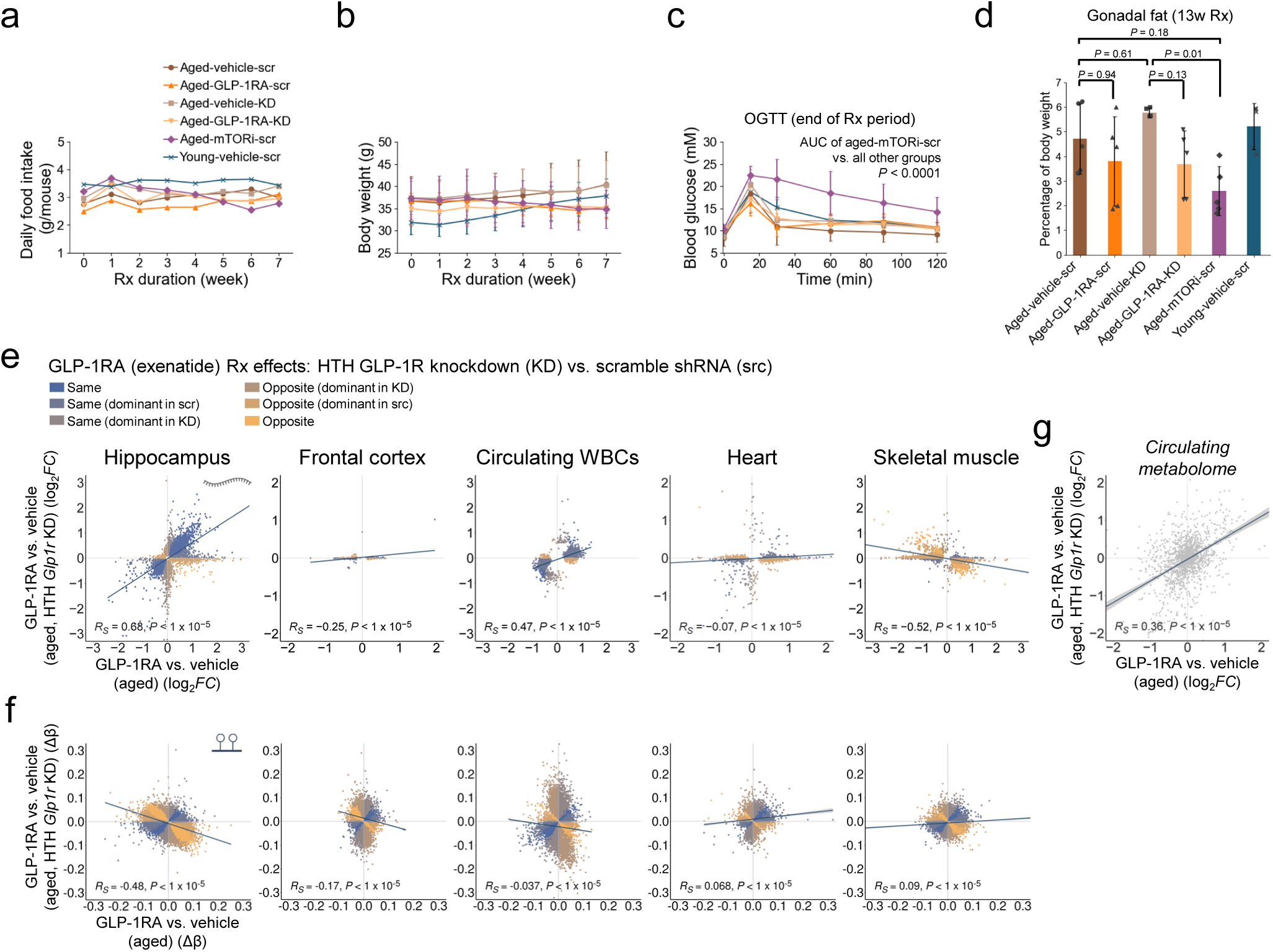
Metabolic changes, age-counteracting multi-omic effects of 13-week GLP-1RA (exenatide) or rapamycin treatment starting at 18 months old, and dependence on hypothalamic GLP-1 receptor. **a**, Longitudinal changes in average daily food intake per mouse for animals in the different experimental groups (monitored weekly). Abbreviations: scr, scramble shRNA group; KD, knockdown group. **b**, Mean (±S.D.) body weight of the animal groups throughout the treatment period (monitored weekly). **c**, Oral glucose tolerance test (OGTT) results for the different experimental groups at the end of 13-week treatment (Rx) period. *P*-value: one-way ANOVA with Holm-Sidak’s post-hoc multiple comparisons test for area under the curve (AUC). **d**, Gonadal fat weight as percentage of body weight in the different experimental groups. *P*-value: one-way ANOVA with Tukey’s post-hoc test for multiple comparisons. **e**, Scatter plots showing transcriptomic changes with exenatide treatment in aged control (scramble shRNA) mice (*x*-axis) vs. hypothalamic (HTH) *Glp1r* knockdown (*y*-axis) mice across the different tissue organs and circulating white blood cells (WBCs). Each dot represents one differentially expressed gene (DEG), color-coded by treatment (Rx) effect categories as shown in the inset (same or opposite Rx effects; dominant in scr or KD: statistically significant in the respective comparison only; no additional label: significant for both comparisons, also see **Methods**). Inset of left panel: symbol for transcript. **f**, Similar to **e**, but showing the results for exenatide treatment-induced differentially methylated loci (DML) covered by the Illumina mouse 285k array (mm285k) across different tissue organs and circulating WBCs. Inset of left panel: symbol for DNA methylation. **g**, Scatter plot showing plasma metabolomic changes with exenatide treatment in aged control (scramble shRNA) mice (*x*-axis) vs. HTH *Glp1r* knockdown (*y*-axis) mice. For each plot in **e**–**g**, the line represents linear fit (with confidence interval (grey) in **g**). The Spearman correlation coefficients (*R_S_*) and associated *P*-values are also shown. Sample sizes for data in all plots: *n* = 5 mice for each experimental group for the various tissues, except *n* = 4 aged exenatide-treated mice for circulating WBCs transcriptomic profiling.

